# Attenuation of ATM signaling by ROS delays replicative senescence at physiological oxygen

**DOI:** 10.1101/2024.06.24.600514

**Authors:** Alexander J. Stuart, Kaori K. Takai, Railia R. Gabbasova, Henry Sanford, Ekaterina V. Vinogradova, Titia de Lange

## Abstract

Replicative senescence, a powerful tumor suppressor pathway, occurs when a few critically-short telomeres activate the DNA damage response (DDR). We show that ATM is the sole DDR kinase responsible for the induction and maintenance of replicative senescence and that ATM inhibition can induce normal cell divisions in senescent cells. Compared to non-physiological atmospheric (∼20%) oxygen, cells grown at physiological (3%) oxygen were more tolerant to critically-short telomeres, explaining their extended replicative lifespan. We show that this tolerance is due to attenuation of the ATM response to double-strand breaks (DSBs) and unprotected telomeres. Our data indicate that the reduced ATM response to DSBs at 3% oxygen is due to increased ROS, which induces disulfide-bridges in ATM, generating crosslinked ATM dimers that do not respond to DSBs. This regulation of cellular lifespan through attenuation of ATM at physiological oxygen has implications for tumor suppression through telomere shortening.

## Introduction

Due to the absence of telomerase activity, the telomeres in most primary human cells undergo progressive shortening. Telomere attrition eventually instigates the Hayflick limit, a stage in the replicative lifespan of human cells where critically-short telomeres induce senescence or apoptosis^1, 2^. This process represents a cell division counting mechanism that culls early-stage tumors when they have depleted their telomere reserve due to an excessive number of cell divisions. Consistent with telomere shortening acting as a tumor suppressor pathway, individuals that are born with abnormally long telomeres are predisposed to a wide variety of cancers, presumably due to accrual of oncogenic changes before the delayed Hayflick limit is reached^3, 4^.

In primary human fibroblasts, senescence is induced when p53 (and to a lesser extent p16) blocks cell cycle progression in response to a DNA damage signal emanating from ∼5 critically-short telomeres^5, 6^. In agreement, cells lacking p53 (or its target p21) readily bypass senescence^7, 8^ and dual inhibition of ATM and ATR kinase signaling with kinase-dead alleles can induce S phase in a subset of senescent BJ fibroblasts^5^. However, since BJ fibroblasts were later shown to express little p16^9^, this reversal of senescence upon dual inhibition of ATM and ATR may have been an exception. It was suggested that critically-short telomeres lack sufficient TRF2^10–12^, the shelterin subunit that represses ATM at telomeres^13^, leading to ATM-dependent senescence. Nonetheless, it remained to be determined whether critically-short telomeres also lack sufficient POT1, the ATR repressor in shelterin, potentially leading to ATR-dependent senescence.

The replicative lifespan of primary human cells is affected by oxygen levels. Cells in most human tissues experience between 0.5 and 8% oxygen, while exposure to the atmospheric (∼20%) oxygen levels used in most tissue culture experiments is limited to cells in the upper trachea^14, 15^. Compared to when grown at atmospheric oxygen levels, cells arrest prematurely at below 0.5% oxygen (hypoxia) or supra-atmospheric (40%) oxygen^16, 17^. When cells are grown at physiological oxygen levels (3% in our experiments), they show a longer replicative lifespan than at 20% oxygen^18–20^. It was proposed that the more rapid senescence at 20% oxygen is partly due to accelerated telomere shortening^18, 21^. However, this proposal is not consistent with the finding that cells cultured for many population doublings (PDs) at low oxygen undergo senescence almost immediately when switched to 20% oxygen whereas the low oxygen controls continue to divide^19^. Furthermore, cells that undergo senescence at low oxygen have slightly shorter, not longer, average telomere lengths^18, 20, 21^ (and see below).

Here we show that the initiation and maintenance of senescence in primary WI38 and MRC5 fibroblasts is solely determined by ATM signaling initiated at short telomeres that lack sufficient TRF2 to repress the activation of this kinase. We establish that the extended replicative lifespan of cells grown at physiological oxygen levels is due to a diminished ATM response to DSBs and damaged or shortened telomeres, and our results implicate increased ROS at 3% oxygen in this attenuation of the ATM response.

## Results

### ATM, not ATR, is required for induction and maintenance of replicative senescence

Two well-established primary human lung-derived fibroblast cell lines, WI38 and MRC5^22, 23^, were used to study replicative senescence. Senescence was observed at population doubling (PD) 54 when WI38 cells were grown in 20% oxygen and at PD62 when cells were cultured at 3% oxygen (Fig. 1A). MRC5 fibroblasts underwent senescence around PD71 and PD77 at 20% and 3% oxygen, respectively (Fig. S1A). The following criteria for senescence were applied to WI38 and MRC5 cells: (1) a maximal PD consistent with the literature^22–24^; (2) fewer than one PD in 10 days; (3) less than 12% of cells showing BrdU incorporation after a 4-day labeling period; and (4) more than 30% of cells staining positive for Senescence Associated β-galactosidase activity^25–28^ (hereafter β-gal) (Fig. 1B-D). In most senescent cell populations between 2 and 7% cells had incorporated BrdU during the 4-day labeling period (Fig. 1C). In many experiments, we included a fifth criterium for senescence: detection of LaminB1 in less than 40% of the cells^25, 27, 29–31^ (Fig. S1B).

**Figure 1.**
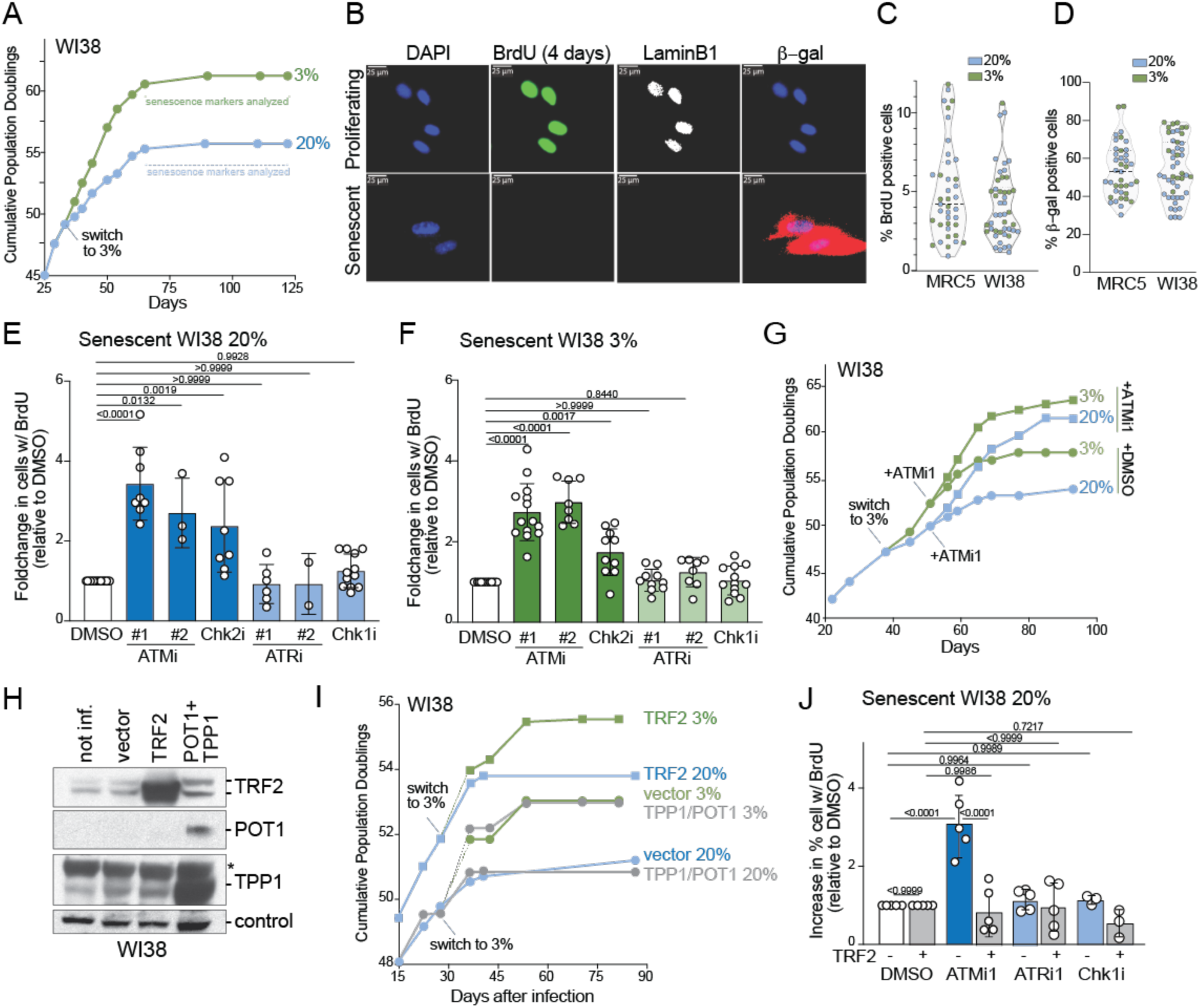
Replicative senescence is due to ATM kinase activation at telomeres lacking sufficient TRF2. **(A)** Representative growth curves of WI38 cells in 20% and 3% oxygen. Horizontal lines denote growth stage at which BrdU incorporation and senescent markers were assayed. **(B)** Examples of BrdU incorporation, LaminB1 staining, and β-gal activity in proliferating and senescent cells. BrdU incorporation was determined by IF after a 4-day labeling period (2 days for proliferating cells). LaminB1 was detected by IF. A fluorescence-based assay was used for β-gal activity. **(C)** Quantification of the % of cells positive for BrdU in senescent MRC5 and WI38 populations determined as assayed in (B). Each data point represents an independent experiment with more than 100 cells scored. **(D)** Quantification of the % of cells positive for β-gal activity in senescent MRC5 and WI38 populations determined as assayed in (B). Each data point represents an independent experiment with more than 100 cells scored. **(E)** Quantification of BrdU incorporation as in (B) after treatment of senescent WI38 cells grown at 20% oxygen with inhibitors of ATM and Chk2 (darker blue) or ATR and Chk1 (lighter blue). Data are represented as relative increase compared to DMSO treatment. Each dot represents an independent experiment with >100 cells scored. n ≥ 3. **(F)** As in (E) but with cells grown to senescence in 3% oxygen. **(G)** Growth curves of WI38 cells with and without long-term ATMi1 treatment at 3% and 20% oxygen. **(H)** Western blots on WI38 cells (PD50) infected with Myc-TRF2 (one infection), FLAG-TPP1+POT1 (two infections) or with the vector control (2 infections). **(I)** Growth curves of WI38 cells as in (H) infected with Myc-TRF2, FLAG-TPP1+POT1, or the empty vector. Cells were first cultured at 20% oxygen and switched to 3% oxygen at the indicated time points or kept at 20% oxygen. **(J)** Quantification of BrdU incorporation as in (E) using Myc-TRF2 overexpressing cells (or the vector control) that had undergone senescence. P values based on One-way ANOVA with Dunnett’s multiple comparisons test for (E) and (F) and Two-way ANOVA with Šídák’s multiple comparisons test for (J). Inhibitors: ATMi1, 670 nM KU-60019; ATMi2, 1.7 µM AZD1390; Chk2i, 2.5 µM BML-277; ATRi1, 1 µM AZD6738; ATRi2, 100 nM AZ20; Chk1i, 50 nM CHIR-124. See also Figure S1.

In senescent WI38 and MRC5 fibroblasts that were incubated with BrdU for 4 days, there was a significant increase in DNA synthesis when cells were treated with inhibitors to the ATM kinase (ATMi) or its downstream effector kinase Chk2 (Chk2i) (Fig. 1E, F; Fig. S1C-E). ATMi and Chk2i induced BrdU incorporation regardless of whether cells had been cultured at 20% or 3% oxygen. In contrast, inhibitors of the ATR kinase (ATRi) or its effector kinase Chk1 (Chk1i) did not induce DNA replication in senescent fibroblasts regardless of the oxygen level (Fig. 1E, F; Fig. S1C-E). Furthermore, addition of ATRi or Chk1i in combination with ATMi did not increase the incorporation of BrdU beyond the effect of ATMi alone (Fig. S1F, G). A combination of ATMi and Chk2i led to a further increase in BrdU incorporation (Fig. S1F, G), possibly due to incomplete ATM inhibition by the ATMi concentration used. A higher ATMi concentration was toxic, perhaps because of an off-target effect (see below Fig. S1I). Live-cell imaging data presented below (Fig. S3) showed no evidence for elevated cell death in cells treated with Chk1i or ATRi, indicating that the absence of increased BrdU incorporation was not due to loss of BrdU-positive cells because they died after their entry into S phase.

Consistent with the finding that ATM is the main kinase that keeps senescent cells from entering S phase, the proliferative lifespan of WI38 and MRC5 cells was significantly extended when cells were cultured in the presence of ATMi at either 3% or 20% oxygen (Fig. 1G; Fig. S1H). Cells treated with ATMi had the same growth rate (0.55 PD/day) as the untreated controls during the early and middle stages of their proliferative lifespan. In ATMi-treated cells that had reached senescence in 3% oxygen, inhibition of the ATR pathway also did not increase the BrdU incorporation (Fig. S1J, K). These data indicate that activation of ATM signaling is the main mechanism by which WI38 and MRC5 fibroblasts arrest in the cell cycle in response to shortened telomeres regardless of the oxygen level.

Consistent with ATM signaling being responsible for entry into senescence, overexpression of TRF2, the shelterin subunit that represses ATM signaling at telomeres^13^, extended the lifespan of WI38 and MRC5 cells at both 3% and 20% oxygen (Fig. 1H, I; Fig. S1L, M). This finding is consistent with the observations that increased TRF2 expression allowed IMR90 cells to better tolerate telomere shortening^12^. In contrast, overexpression of the TPP1/POT1 heterodimer in shelterin, which represses ATR but not ATM signaling at telomeres^13^, had no effect and neither did overexpression of the TRF2-interacting partner Rap1 (Fig. 1H, I; Fig. S1L, M). As expected, cells that had reached senescence despite overexpression of TRF2 did not respond to ATMi with an increase in BrdU incorporation (Fig. 1J; Fig. S1N, O). They also did not increase their BrdU incorporation in the presence of ATRi (Fig. 1J), making it unlikely that these cells underwent senescence due to a deficiency of POT1. In contrast, vector-infected control cells still showed an increase in BrdU incorporation in response to ATMi. Based on these data, we conclude that replicative senescence is initiated by critically short telomeres that induce ATM kinase signaling because they lack sufficient TRF2. In agreement, the shortest telomeres in senescent cells exhibit diminished TRF2 IF signals^10^.

### ATMi can reverse replicative senescence

Since ATM inhibitors reproducibly induced entry into S phase in a fraction of senescent cells, we examined the fate of these cells using live-cell imaging. Prior to entry into senescence, MRC5 cells were infected with a fluorescent ubiquitination-based cell cycle indicator (FUCCI^32^) vector to identify cells in G1, S, and G2 (Fig. 2A). Cells were automatically tracked^33^ in images captured every 30 min during 44-97 h imaging sessions (Videos S1-4), and their FUCCI signals were processed to generate tracks in which S/G2 is indicated with green and G1 in grey (Fig. 2A; Fig. S2B-G). Cells undergoing mitosis (indicated with yellow) were verified by visual inspection. Approximately half of pre-senescent MRC5 cells (at ∼PD66) entered S phase and divided once or twice in a 44.5-h time window (Fig. 2B). Most cell divisions were normal, although there were few (<12%) aberrant cell cycle events which is expected for a cell population nearing senescence. In contrast, senescent MRC5 cells (imaged at PD78) showed minimal entry into S phase (11% of cells), infrequent normal cell divisions (<4% of all cells), and many cells that entered S phase but did not proceed through a normal cell division (Fig. 2E). This low – but consistent – frequency of aberrant cell cycle events is in agreement with reports of bi-nucleated and tetraploid cells in senescent and near-senescent cultures^34–37^. Inhibition of the kinase activity of ATR or Chk1 had no effect on senescent cells (Fig. S3A, B). In contrast, ATMi not only induced frequent entry into S phase (∼25% of cells) but also instigated complete and apparently normal cell divisions after resumption of the cell cycle (Fig. 2D, E; Videos S3, S4). In some cases, two or three consecutive cell divisions were recorded in less than 3 days of imaging (Fig. 2E-G).

**Figure 2.**
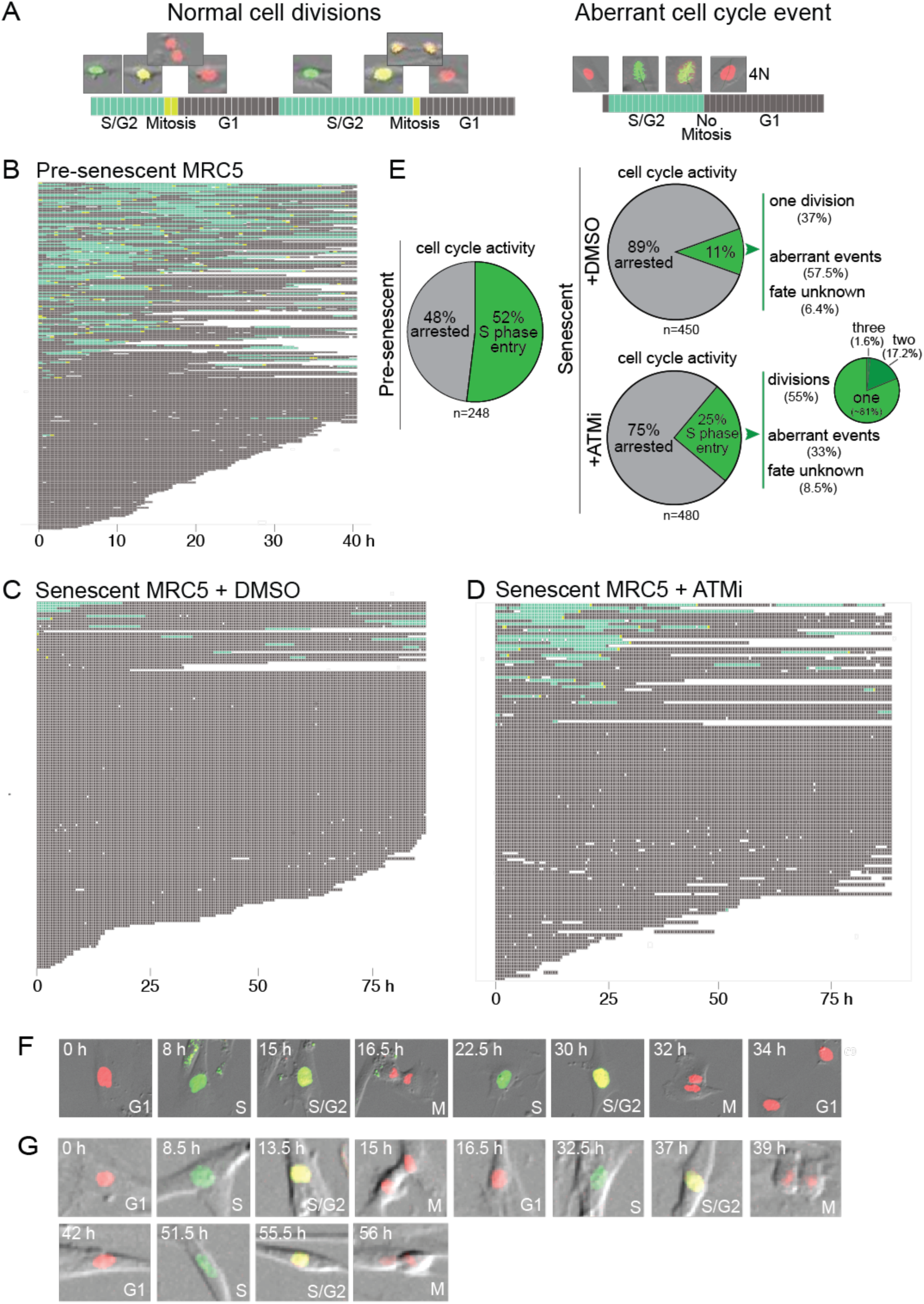
ATM inhibition induces cell divisions in senescent MRC5 cells. **(A)** Left: Representative live-cell FUCCI imaging (30 min intervals) track and example images of pre-senescent MRC5 cells undergoing normal cell divisions. Cell cycle stages represented by colors in the tracks: green, S/G2; yellow, mitosis; grey, G1. Right: Representative track and images of an aberrant cell cycle event in an ATMi-treated senescent cell that by-passed mitosis leading to a 4N G1 cell. **(B)** All recorded cell tracks of pre-senescent MRC5 cells. Rows represent individual tracked cells (n=248). Imaging times indicated below. White indicate gaps due to uninterpretable images or cells leaving the field. At least 74% of cycling cells undergo at least one normal division and 14% had an unknown fate due to the end of the imaging session or the cell exiting the field. **(C)** As in (B) but for senescent MRC5 cells (n=144). Representative of 3 replicates. **(D)** As in (B) but for senescent MRC5 cells treated with ATMi1 (670 nM) (n=122). Representative of 3 replicates. **(E)** Comparison of cell cycle activity in pre-senescent MRC5 cells and senescent MRC5 cells treated with DMSO or ATMi1. Data from 3 biological replicates with total numbers of tracked cells indicated. Smaller pie chart (right) representing cells undergoing at least one division. **(F) and (G)** Images of cells undergoing multiple divisions in two independent ATMi-treated senescent MRC5 cell cultures. See also Figures S1, S3 and Videos S1-4.

These data reveal that ATMi can reverse the senescent state in at least a subset of the cells. We note that most of the MRC5 cells used here had been in senescence for approximately one month. It remains to be seen whether ATMi has the same effect on cells that have been senescent for longer time periods when the Senescence Associated Secretory Pathway (SASP) and its paracrine effects are likely to generate a state that is not sensitive to ATMi^38, 39^.

### Attenuation of ATM DNA damage signaling at 3% oxygen

The accelerated replicative senescence at 20% oxygen could be explained if the cells have a higher rate of telomere attrition compared to their counterparts grown at 3% oxygen. This explanation is not consistent with our findings (Fig. 3A) and those of others^19^ that pre-senescent cells that had been extensively propagated at physiological oxygen can show sudden senescence when they are switched to 20% oxygen. In our experiments, cells were grown in 3% oxygen to a PD where they would have reached senescence when growing at 20% oxygen. When such cells were transitioned from 3% to 20% oxygen, they stopped growing almost immediately (Fig. 3A), indicating that the cells responded to the same telomere length differently at 20% oxygen. Furthermore, when cells underwent this transition in the presence of ATMi, they continued growing (Fig. 3A). These observations suggest that cells display a sudden increase in the ATM response to short telomeres when they are transferred to 20% oxygen, as opposed to a change in telomere attrition. Indeed, in agreement with prior data^18, 20, 21^, our genomic blotting and Universal Single Telomere Length Analysis (USTELA^40^) in which telomeric fragments are amplified to ascertain the lengths of individual telomeres, showed that telomeres of senescent WI38 and MRC5 cells were not shorter when the cells were cultured at 20% oxygen (Fig. 3B; Fig. S4A,B). In fact, the burden of very short telomeres detected with USTELA appeared significantly higher in cells that had senesced at 3% oxygen (Fig. 3B, C; Fig. S4B, C), indicating that very short telomeres are better tolerated at low oxygen. The greater tolerance for short telomeres at low oxygen was not due to obvious changes in expression of TRF2, ATM, Mre11, or Chk2 (Fig. 3D).

**Figure 3.**
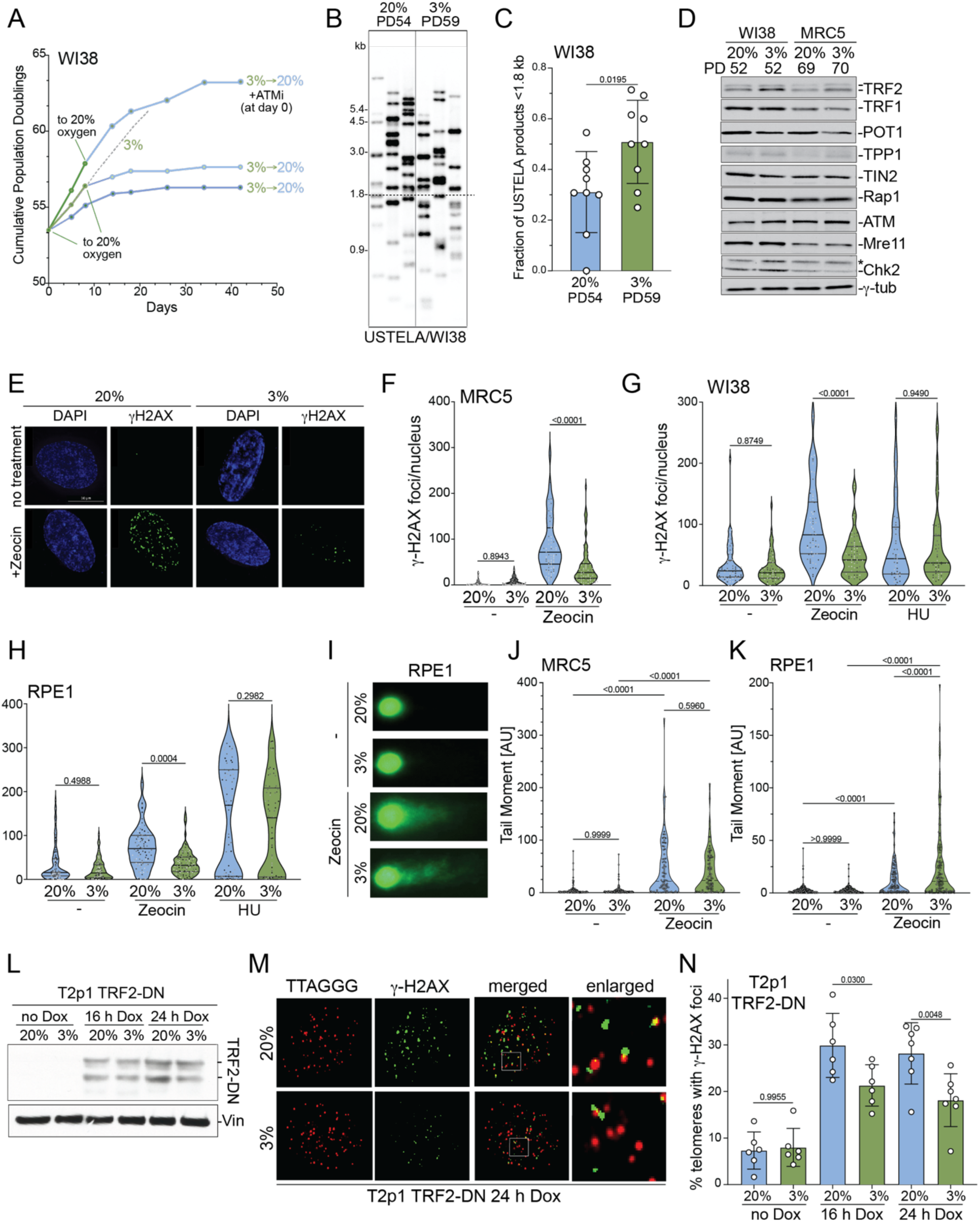
Reduced ATM response to DSBs and unprotected telomeres at 3% oxygen. **(A)** Growth curve of WI38 cells grown at 3% oxygen and switched to 20% oxygen as indicated, with or without 670 nM ATMi1. The switch from 3 to 20% oxygen was at a PD close to the final PD in a 20% oxygen-only culture (see Fig. 1F). Dotted line represents growth trajectory of untreated cells at 3% oxygen from Fig. 1F. **(B)** USTELA products from WI38 cells grown to senescence at high and low oxygen. The three lanes for each sample represent three independent USTELA PCRs. **(C)** Quantification of fraction of USTELA products shorter than 1.8 kb for each PCR as in (B) (n=9). **(D)** Western blots for the indicated proteins from WI38 and MRC5 cells grown at low or high oxygen. **(E)** Representative images of the DNA damage response in MRC5 cells grown at 20% or 3% oxygen. Cells were treated with zeocin (or not) and stained for DNA (DAPI, blue) or γ-H2AX (IF, green). **(F -H)** Quantification of γ-H2AX foci in proliferating MRC5, WI38, or RPE1 cells treated with zeocin (50 µg/mL, 2.5 h) or HU (2 mM, 2.5 h). n≥3 with 20-25 cells quantified per experiment. (**I)** Representative image of comet assays on RPE1 cells treated as indicated. DNA stained with 1X SYBR Gold. **(J and K)** Quantification of comet assay tail moment in proliferating MRC5 and RPE1 cells at the indicated oxygen levels with and without zeocin treatment as in (F) and (H). **(L)** Western blot for TRF2-DN induction 16 and 24 h after addition of 1 µg/ml of doxycyclin in T2p1 cells proliferating at 3% or 20% oxygen. The lower band is a degradation product. Vinculin (vin) serves as a loading control. **(M)** IF-FISH detecting TRF2-DN induced γ-H2AX foci at telomeres (TIFs) in T2p1 at 3% and 20% oxygen. **(N)** Quantification of TIFs detected as in (M). Data is from 6 independent experiments with 20-30 nuclei quantified in each. P values were determined as follows: (C) unpaired t-test; (F-H) and (N) One Way ANOVA with Šídák’s multiple comparison; (J) and (K) One Way ANOVA with Tukey’s multiple comparison. See also Figure S4.

Given the reduced ATM response to short telomeres at 3% oxygen, we tested whether the ATM kinase also responded less to DSBs at 3% vs. 20% oxygen. Strikingly, the formation of γ-H2AX foci in response to zeocin-induced DSBs, a quantifiable proxy for ATM activation, was 2.25-fold and 2.1-fold lower at 3% oxygen in MRC5 and WI38 cells, respectively (Fig. 3E-G). Similarly, the response at ATM to DSBs induced by etoposide was reduced at 3% oxygen (Fig. S4D). In contrast, oxygen levels did not affect the response to HU, which induces replication stress that activates the ATR kinase (Fig. 3G). Like the primary human fibroblasts, RPE1 cells showed a 2.1-fold lowered response to DSBs, but not to replication stress, at 3% vs. 20% oxygen (Fig. 3H). Comet assays to detect DSBs showed that zeocin induced the same level of DNA damage at 3% and 20% oxygen (Fig. 3I-K). The attenuation of the response of ATM to DSBs was also observed in a previously characterized RPE1-based system where induction of a dominant-negative allele of TRF2 activates ATM kinase signaling at dysfunctional telomeres^11,41^ (Fig. 3L-N). These data suggest that the ATM kinase responds to DSBs and dysfunctional telomeres more robustly at 20% oxygen, explaining why senescence is accelerated at high oxygen.

As HIF-1α is induced at low oxygen and promotes cell survival^42–46^, it was prudent to determine whether HIF-1α is responsible for the attenuated response of ATM to DSBs and critically short telomeres at low oxygen. An inducible shRNA that reduced HIF-1α expression detectable by western blotting and IF did not affect the induction of γ-H2AX foci in response to zeocin at 3% or 20% oxygen (Fig. S4E-G). As expected, MRC5 cells treated with an inhibitor of HIF-1α translation (HIF-1αi) had a reduced proliferative lifespan at low oxygen^41^ (Fig. S4H). However, consistent with ATM being upstream of HIF-1α^43, 47^, this lifespan reduction was not due to increased ATM signaling since addition of ATMi did not increase the lifespan of HIF-1α treated cells (Fig. S4H).

### Increased ROS at low oxygen diminishes the ATM response to DNA damage

ATM dimers can become covalently linked through cysteine disulfide-bridges which form in response to H_2_O_2_ generated by other reactive oxygen species (ROS), including superoxide^48, 49^. These crosslinked ATM dimers are active and target factors that increase ROS tolerance but do not induce a cell cycle checkpoint in untreated, physiologically relevant settings^16, 43, 50, 51^. Importantly, ROS-induced crosslinked ATM dimers cannot be activated at DNA breaks. At DSBs, ATM is activated when MRN bound to DNA ends converts ATM into active monomers which enforce the cell cycle checkpoint by phosphorylating Chk2 and p53^50^ (see Fig. 4N below). This formation of active ATM monomers at DSBs is not possible when the ATM dimers are held together by disulfide bridges.

**Figure 4.**
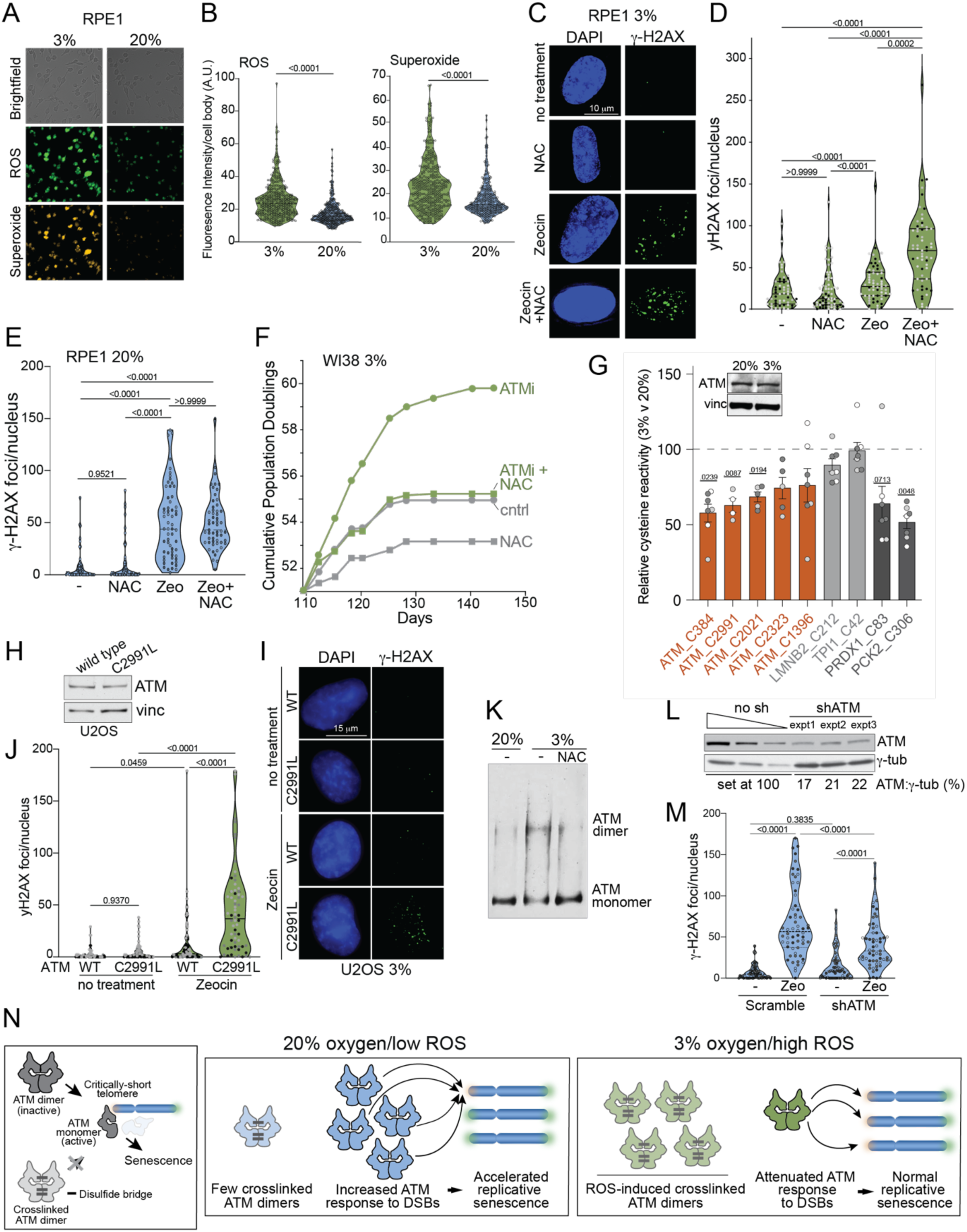
ROS-induced ATM crosslinking attenuates the DSB response at 3% oxygen. **(A)** Representative images of fluorescence staining for ROS and Superoxide in RPE1 cells growing at 20% and 3% oxygen. **(B)** Quantification of ROS and Superoxide in RPE1 cells at 3 and 20% oxygen determined as in (A). **(C)** Representative γ-H2AX IF in RPE1 cells grown at 3% oxygen after treatment with zeocin (2.5 h, 50 µg/mL), NAC (2 days, 5 mM), or both (2 days with NAC and zeocin added 2.5 h before fixation). **(D)** Quantification of γ-H2AX foci formation in RPE1 cells treated as in (C). n = 3, at least 20 nuclei per experiment. **(E)** Quantification of γ-H2AX foci formation in RPE1 cells grown at 20% oxygen after treatment with zeocin (2h, 25 µg/mL), NAC (2 days, 5 mM), or both as in (C). n = 3 with 15-20 cells per experiment **(F)** Growth curves of WI38 cells grown at 3% oxygen treated with ATMi1 (670 nM) and NAC (5 mM) as indicated. Compounds added at day 109. **(G)** Relative reactivity for the indicated cysteine residues in ATM and control proteins. Values represent the ratio of the signal intensity for peptides containing the denoted cysteine residue in samples from cells grown at 3% vs. cells grown at 20% oxygen. Error bars represent standard error of the mean (SEM). For each cysteine, sets of data points with identical shading (light grey, dark grey, white) represent technical replicates, whereas the different shadings represent biological replicates. Statistics were based on the average value for three biological replicates determined from the means of the technical replicates. Inset: immunoblot to determine the expression level of ATM at the 20% and 3% oxygen. Vinculin (vinc) serves as a loading control. **(H)** Immunoblot for ATM in U2OS cells expressing the wildtype or mutant version. Vinculin serves as the loading control. **(I)** Representative images of γ-H2AX foci in U2OS cells expressing either wild type (WT) ATM or the ATM-C2991L mutant grown at 3% oxygen and treated (or not) with 50 µg/ml zeocin for 1 h. **(J)** Quantification of γ-H2AX foci formation in U2OS cells as in (J). n = 3, at least 15 cells per experiment. **(K)** Detection of ATM dimers and monomers in immunoblots of native gel fractionation of RPE1 cell extracts. RPE1 cells were grown at 20% oxygen, 3% oxygen, or 3% oxygen and treated with 5 mM NAC. **(L)** Immunoblot to determine the reduction in ATM expression upon shRNA treatment. Extracts from control RPE1 cells without shRNA treatment were diluted 1:2 and 1:4 in the second and third lane to ensure that the signals were in the linear range. Ratios of ATM signal to γ-tubulin (γ-tub) signal were quantified and expressed as a % of the ratio in the control cells. **(M)** Quantification of γ-H2AX foci formation after zeocin treatment in RPE1 cells treated with the ATM shRNA as in (K). n = 3, at least 20 nuclei per experiment. Cells treated (or not) with 50 µg/mL zeocin, 2.5 hrs. **(N)** Model for the effect of oxygen on ATM activation at DSBs and critically-short telomeres. At physiological (3%) oxygen, higher ROS induces disulfide-bridges in ATM creating crosslinked ATM dimers that do not respond to DNA damage. The diminished pool of ATM that can respond to DSBs allows cells to tolerate critically-short telomeres better and divide more often. At 20% oxygen, there is less ROS-induced ATM crosslinking, resulting in a greater pool of ATM that can respond to DNA damage. This ‘hyperactive’ ATM pool responds more robustly to critically-short telomeres, resulting in accelerated replicative senescence at normoxia. P values determined by unpaired t-test for (B), Two-Way ANOVA with Tukey’s multiple comparison for (D), and Ordinary One-Way ANOVA with Tukey’s multiple comparison for (E), (J) and (M). The p value for the diminished reactivity of the cysteines in (G) was determined by One Sample t and Wilcoxin test based on the averages of median values from technical replicates. The data on the cysteines in mitochondrial proteins and the controls were treated similarly. See also Figure S4.

Paradoxically, low oxygen conditions induce reactive oxygen species (ROS) to a greater extent than at 20% oxygen through a mechanism that is not fully understood^25, 42, 45, 52, 53^. We confirmed the increase in ROS and superoxide production in RPE1, MRC5, and WI38 cells at 3% oxygen compared to 20% oxygen (Fig. 4A,B; Fig. S4J,K). Furthermore, as expected, overexpression of TRF2 did not ablate the increase of ROS in 3% oxygen conditions (Fig. S4L). This increase in ROS at low oxygen might induce ATM dimer crosslinking, explaining the diminished ATM response to DNA damage.

To test whether the ATM response to DSBs and critically-short telomeres is diminished due to ROS-induced ATM dimer crosslinking at low oxygen, we used N-acetyl-cysteine (NAC) which removes the crosslinks from ATM^48^. At low oxygen, NAC increased the frequency of γ-H2AX foci in response to zeocin (Fig. 4C, D), consistent with NAC minimizing crosslinked ATM dimers that do not respond to DSBs. At 20% oxygen, the effect of NAC on the DNA damage response was not significant (Fig. 4E). Furthermore, NAC reduced the proliferative lifespan of WI38 and MRC5 cells grown at low oxygen and this effect was in part due to increased ATM activation since treatment with ATMi extended the lifespan of NAC-treated cells (Fig. 4F, S4I). Similarly, treatment with disulfide bond-reducer Tris(2-carboxyethyl)phosphine hydrochloride (TCEP) increased γ-H2AX foci formation upon zeocin treatment and counteracted the proliferative benefit of ATMi (Fig. S4M,N). These findings are consistent with the observation that resveratrol treatment, which reduces ROS, increases the ATM response to bleomycin-induced DSBs^54^.

To monitor the effect of 3% oxygen on the formation of disulfide bridges in ATM directly, we used chemical proteomics with sample multiplexing (TMT-ABPP) to directly compare the reactivity of cysteines with an iodoacetamide-desthiobiotin probe on RPE1 cells grown at 20% and 3% oxygen. In this method, reactive cysteines are covalently linked to an iodoacetamide tag containing a desthiobiotin enrichment handle, allowing the quantification of changes in cysteine reactivity by mass spectrometry^55, 56^. Despite equal expression of ATM determined by immunoblotting, three of the five cysteine residues that were quantified showed significantly lower signal intensity at low oxygen (Fig. 4G, Table S1). Among these was residue C2991, which was previously shown to form a disulfide-bridge upon exposure to ROS^5^. As a positive control, a cysteine in the mitochondrial PCK2 protein, which is in proximity to the presumed site of ROS generation at 3% oxygen showed significantly reduced signal intensity at low oxygen (Fig. 4G, Table S1). A second positive control, C83 in PRDX1, which is known to dimerize through disulfide bridges in response to ROS^57^, also appeared less reactive although this was not statistically significant. In contrast, the accessibility of two negative control cysteines (LaminB2 C212 and TPI1 C42) was unchanged at 3% oxygen (Fig. 4G, Table S1). In addition, there were a number of cysteine-containing peptides derived from other proteins that showed reduced signal intensity, raising the possibility that the higher ROS at 3% oxygen may be affecting the function of additional pathways. These effects will be pursued in the future.

We determined the role of disulfide bridge formation in the ATM DDR using U2OS cells that overexpress either wild type ATM or the C2991L mutant of ATM in which C2991 cannot form a disulfide bond^48^. The C2991L mutant of ATM was previously shown to be hampered in its response to ROS^48^. In cells grown at 3% oxygen, the formation of zeocin-induced γ-H2AX foci was much greater in cells expressing the ATM C2991L mutant than in cells expressing wild type ATM (Fig. 4H-J).

We also used non-denaturing gels and immunoblotting to monitor for the presence of the larger molecular weight species of ATM that represents the crosslinked ATM dimer^48, 57^ . This slower migrating form of ATM, presumably representing the dimer, was detected in cells grown at 3% oxygen but less abundant in cells grown at 20% oxygen and in cells grown at 3% oxygen treated with NAC (Fig. 4K).

Finally, we used an shRNA to the ATM kinase to determine whether partial reduction of the available ATM protein leads to a diminished level of γ-H2AX foci formation after zeocin treatment (Fig. 4L, M). Quantitative western blotting showed that the ATM shRNA reduced the abundance of ATM protein by ∼80%. Under these conditions, the cells showed about ∼60% fewer γ-H2AX foci after zeocin treatment (Fig. 4L, M). This result is consistent with the assumption in the model in Fig. 4N that the levels of ATM protein that can respond to DSBs are limiting for the response to DSBs and critically-shortened telomeres.

Collectively, these findings support the model in which ROS-induced disulfide bridges reduce the ability of ATM to respond to DSBs and critically-short telomeres at physiological oxygen (Fig. 4N).

## Discussion

The results presented here establish that replicative senescence occurs when ATM signaling is activated at critically-short telomeres lacking sufficient TRF2 (Fig. 4N). We imagine that the critically-short telomeres are no longer capable of recruiting sufficient TRF2 to maintain the protective t-loop configuration^58^, allowing MRN to recognize the telomere end and activate ATM. Direct demonstration of t-loop loss at critically-short telomeres is currently not feasible due to their predicted low abundance in senescent cells^6^ and the limitations of the t-loop assay^58^ . Furthermore, the exact length of the TTAGGG repeat region required to avoid ATM activation at telomeres is currently unknown. These questions merit attention in future studies.

Our data also explain why primary human cells have an extended lifespan at low oxygen, or rather, why cells cultured at atmospheric oxygen show accelerated senescence. We show that at 20% oxygen, a condition generally used for studying the DNA damage response, the ATM kinase has an increased ability to respond to DSBs. As a result, cells have a reduced tolerance for short telomeres at 20% oxygen and senescence is induced prematurely (Fig. 4N). We propose that at physiological oxygen, the increased ROS crosslinks more ATM dimers, resulting in a dampened ATM response to DSBs, a greater tolerance of short telomeres, and a longer lifespan (Fig. 4N).

The finding that oxygen levels affect the ability of ATM to respond to DNA damage and critically-short telomeres may have implications for cancer radiotherapy. Lowered ATM activity is known to enhance the efficacy of radiotherapy but also leads to an increase in radiation toxicity^59^. In this context it is prudent to consider how the reduced ATM response to DSBs at lower oxygen levels may affect the therapeutic outcomes and side effects of radiotherapy.

It is now clear that telomere shortening prevents the development of a wide variety of human cancers and it is assumed that this tumor suppressor pathway acts by erecting a proliferative barrier through the induction of senescence or apoptosis. In agreement with ATM as the dominant player in the induction and maintenance of replicative senescence, ATM mutations were frequent in clonal hematopoiesis in patients with telomere biology disorders ^60^.

In frank tumors that have overcome the telomere tumor suppressor pathway through the activation of telomerase, telomeres are often maintained at a relatively short length^61^. For instance, average telomere lengths of cervical and endometrial carcinomas are reported to be less than 3 kb, predicting that these cancers carry a significant burden of telomeres at or near the critical telomere length^61^. Our findings predict that telomere shortening is better tolerated under the low oxygen conditions that are common during tumorigenesis. Therefore, the number of telomeres that need to shorten beyond the minimal length required for ATM repression is likely to be greater than the 4-5 telomeres estimated to initiate the Hayflick limit at 20% oxygen^6^. The greater tolerance for critically-short telomeres at the low oxygen likely to be experienced by cells that make up these cancers explains why there is no selection for greater telomere extension by telomerase. If this reasoning is correct, ATM kinase agonists, relevant disulfide reducers (i.e., TRX1 ^57^), or treatments that reduce ROS could have a therapeutic utility in cancers with very short telomeres.

## Limitations of Study

In this study, we show that ROS-induced disulfide crosslinks are responsible for the reduced ability of ATM to respond to DSBs and critically-short telomeres at physiological oxygen. However, we have not elucidated how ROS is increased at low oxygen and with the exception of C2991, we have not determined which of the many cysteines in ATM are involved in the crosslinking of ATM dimers. We also have not addressed additional questions, including at what length telomeres lose their protection at 3% and 20% oxygen, whether this minimal length is dependent on oxygen conditions, and how many critically short telomeres are needed to induce cell cycle arrest at 3%.

## Resource availability

All data are available in the main text or the supplementary materials. Raw mass spectrometry data will be uploaded to PRIDE. Rscript code is available online at https://github.com/astu17/Trackmate-FUCCI-Code.git.

## Acknowledgments

We thank Gerald Shadel and Yichong Zhang for advice regarding the analysis of ATM on non-denaturing gels. We thank Kivanc Birsoy and Mike Kastan for advice and Bill Kaelin for discussion and the shHIF-1α plasmid. John Maciejowski and the Rockefeller Bio-imaging Resource Center members Crystina Pygaki and Calos Rico are thanked for help with live-cell imaging. Erica Wolin and Jamie Phipps are thanked for help with coding. Agata Smogorzewska, Cayla Broton, and Nicky Blobel are thanked for help with the comet assay. We are grateful to John Sedivy who provided helpful comments on the manuscript. We thank Logan Myler for his advice throughout the project. The U2OS cells were provided by Tanya Paull. We thank all current and past members of the de Lange lab who provided advice during this project. This work was supported by grants from the NIH to T.d.L. (R35CA210036 and R01AG016642). E.V.V. is supported by Rockefeller University start-up funds, Robertson Foundation, The Achelis and Bodman Foundation, and Searle Scholar Program. E.V.V. and H.S. were supported by the Stavros Niarchos Foundation (SNF) as part of its grant to the SNF Institute for Global Infectious Disease Research at The Rockefeller University. Some analyses were performed on computational resources from the Rockefeller University High Performance Computing Resource Center, RRID: SCR_025889.

## Author contributions

Conceptualization, methodology, investigation, data visualization, and manuscript editing by TdL and AJS. Project administration, supervision, funding acquisition, and original written draft by TdL. KKT performed Western blots and experiments with shRNAs to ATM. Cysteine reactivity assays were performed by AJS, RR, HS, and EVV.

## Supplemental Item Titles and Legends

**Figure S1.**
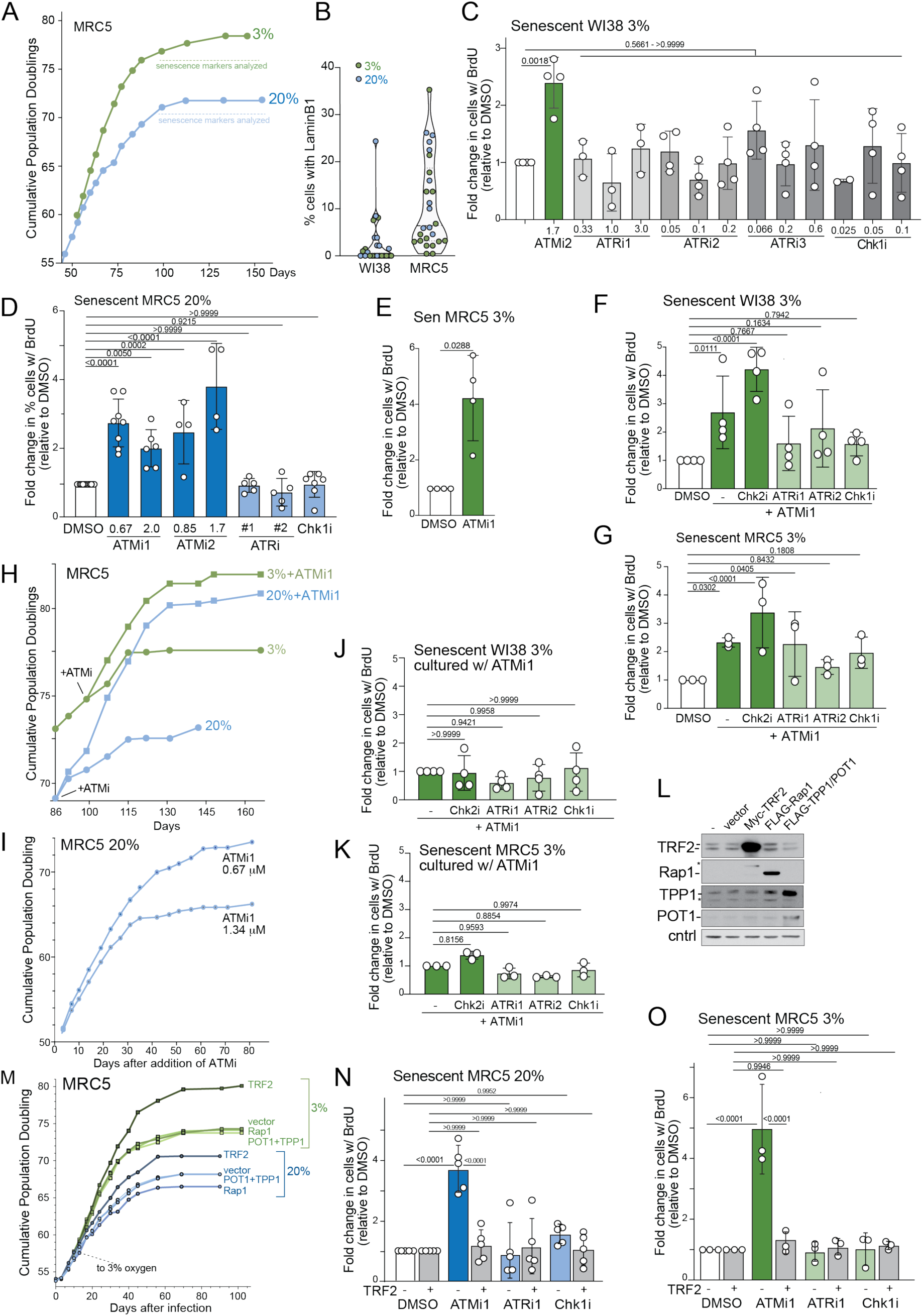
Additional evidence that replicative senescence is due to ATM kinase activation at telomeres lacking sufficient TRF2, related to Figure 1. **(A)** Representative growth curves of MRC5 cells grown at 20% and 3% oxygen as in Figure 1A for WI38 cells. **(B)** Quantification of fraction of cells expressing LaminB1 in senescent WI38 and MRC5 cultures. **(C)** Additional data on the effect of ATMi, ATRi, and Chk1i on the BrdU incorporation in WI38 cells grown to senescence in 3% oxygen analyzed as in Figure 1E. **(D)** and **(E)** Effect of the indicated concentrations of ATMi, ATRi, Chk2i, and Chk1i on the BrdU incorporation in MRC5 cells grown to senescence in 20% or 3% oxygen analyzed as in Figure 1E and 1F. **(F)** and **(G)** Effect of combining ATMi with Chk2i, ATRi, or Chki on BrdU incorporation in the indicated cells grown to senescence at 3% oxygen. **(H)** MRC5 growth curves with and without long-term ATMi1 (KU-60019, 670 nM) treatment at 3% and 20% oxygen. **(I)** Effect of ATMi1 at 670 nM and 1.34 µM on MRC5 cells grown at 20% oxygen. **(J)** and **(K)** Effect of combinations of ATMi with Chk2i, ATRi, or Chk1i on the BrdU incorporation of WI38 (J) or MRC5 (K) cells that had grown to senescence in the presence of ATMi. Experiments represent parallel growth, treatment, and quantification with the cells in (F) and (G). **(L)** Western blots for the indicated shelterin subunits in uninfected MRC5 cells or cells infected with TRF2, Rap1, TPP1/POT1, or a vector control as indicated. **(M)** Growth curves of MRC5 cells in (K). **(N)** and **(O)** Quantification of BrdU incorporation in Myc-TRF2 overexpressing cells (or the vector control) that had undergone senescence. Cell infected with Myc-TRF2 or the empty vector were cultured into senescence and then treated with inhibitors of ATM, ATR or Chk1 as indicated during a 4-day incubation with BrdU. P values determined by Ordinary One-way ANOVA with Dunnett’s multiple comparisons test for (C), (D), and (E); Two-way ANOVA with Dunnett’s multiple comparisons test for (G) and (K); Two-way ANOVA with Šídák’s multiple comparisons test for (F), (J), (N) and (O). ATMi, KU-60019; ATMi2, AZD1390; ATRi1. AZD6738; ATRi2, AZ20; ATRi3, M4344; Chk1i, CHIR-124.

**Figure S2.**
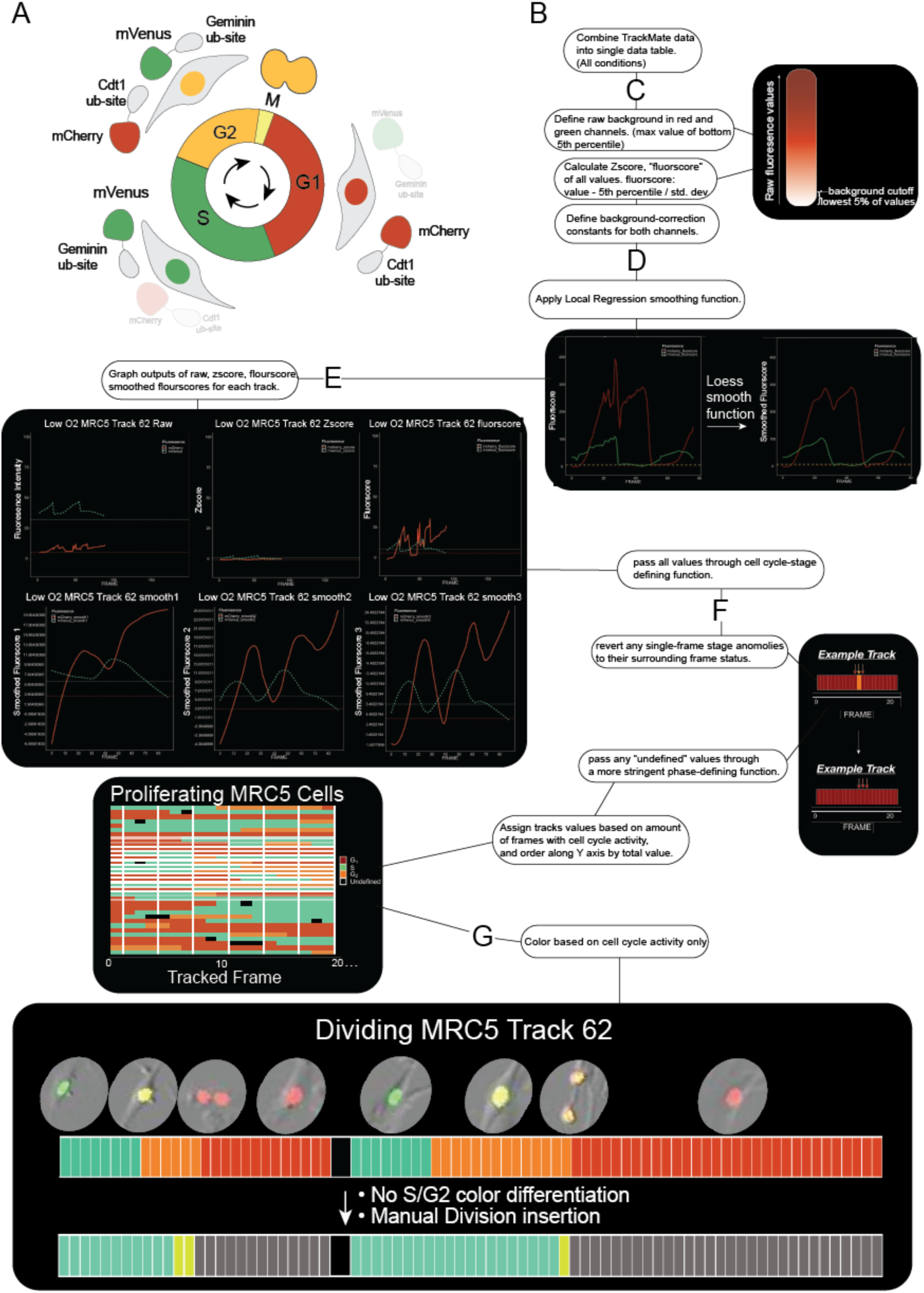
Workflow for analysis of FUCCI live-cell imaging, related to Figure 2. **(A)** Schematic for FUCCI with cell cycle-controlled degrons of Cdt1 and Geminin fused to mCherry and mVenus, respectively. **(B)** Rscript pipeline for largescale Trackmate data analysis. **(C)** Background for both channels defined by the 5^th^ percentile raw value. **(D)** Local regression smoothing function applied to data, which filters out anomalies arising from z-plane changes, debris, etc. **(E)** Graphical output of data for each individual track. Representative section of larger data set. **(G)** Tracks visualized based on cell cycle activity.

**Figure S3.**
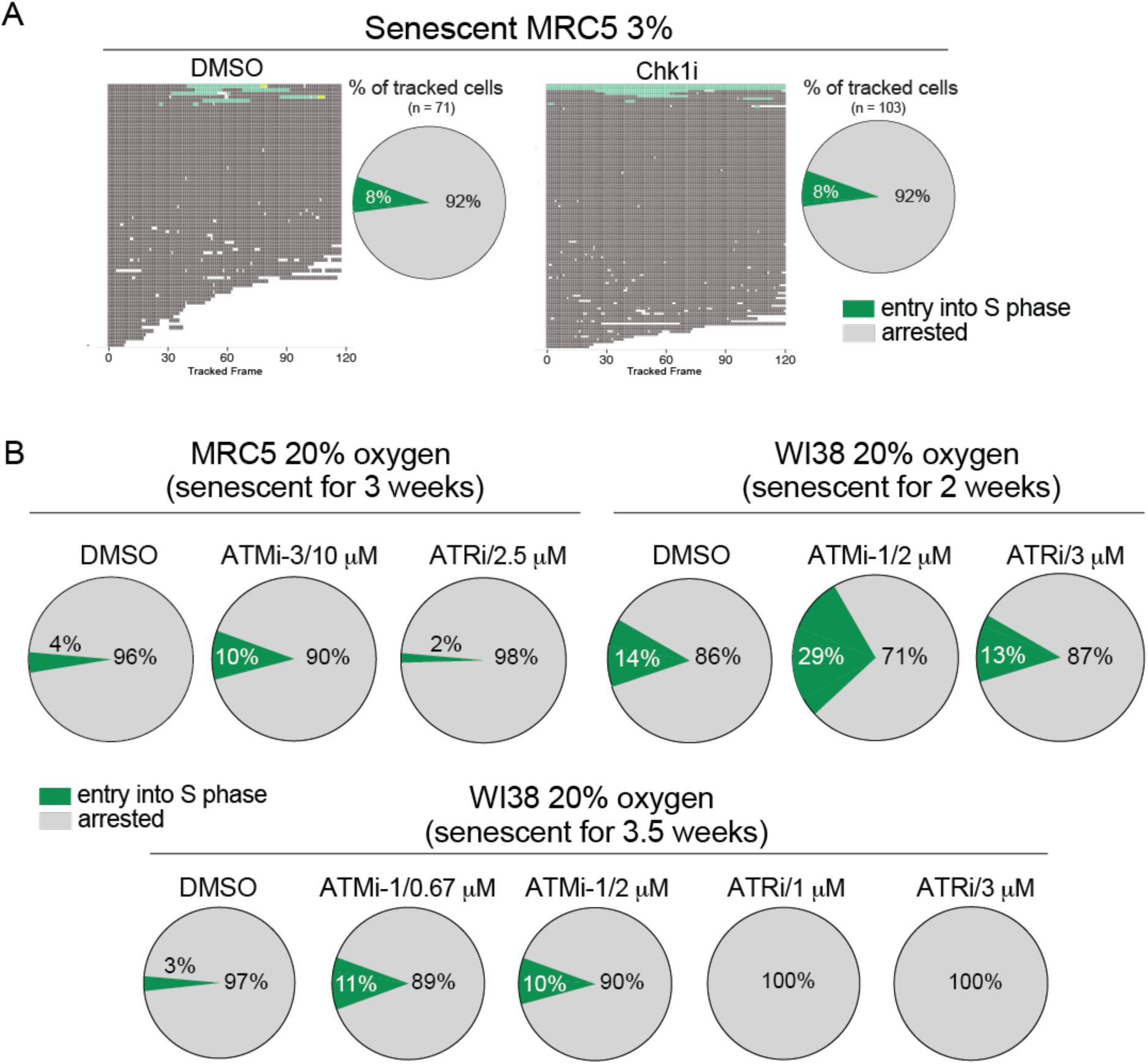
Additional evidence for the reversal of senescence by ATMi but not ATRi, related to Figure 2. **(A)** Imaging tracks and summary of cell cycle stages of MRC5 cells grown to senescence in 3% oxygen treated with Chk1i (CHIR-124, 50 nM) as in Figure 2. Data on control senescent MRC5 cells from Figure 2C and E is shown for comparison. **(B)** Top, left: Cell cycle activity (as in Figure 2E) of MRC5 cells grown to senescence in 20% oxygen based on tracking 150 cells from 97-h imaging sessions. ATMi-3, KU-55933 at 10 µM. Top, right: Cell cycle activity of WI38 cells grown to senescence in 20% oxygen. Experiment conducted after 2 weeks of reaching less than one PD per 10 days. ≥28 cells from a 98-h imaging session. Bottom: Experiment conducted 1.5 weeks later compared to WI38 cells shown above tracking 34 cells in a 94.5-h imaging session.

**Figure S4.**
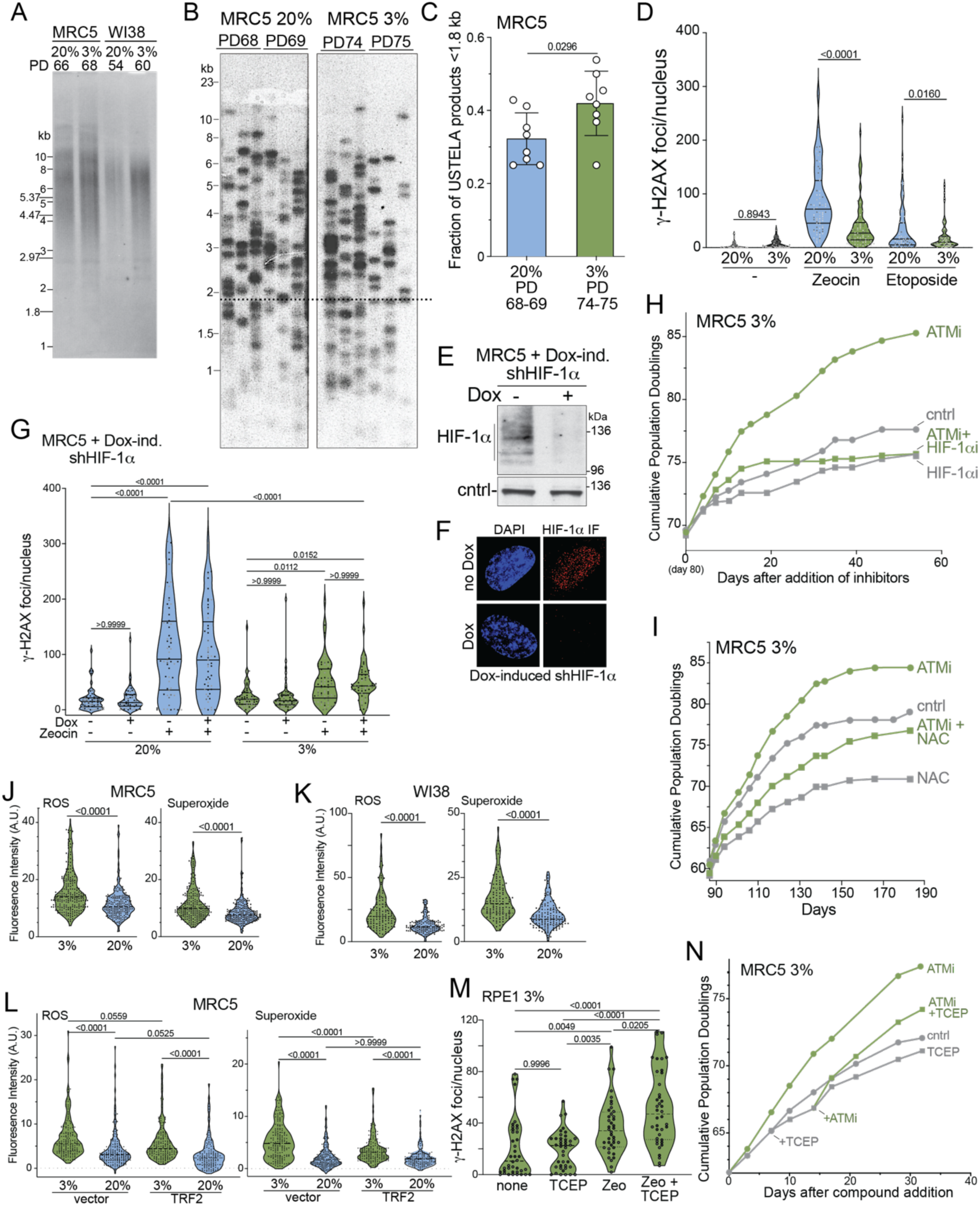
Greater ATM response to DSBs, not telomere shortening rates explains early senescence at 20% oxygen, related to Figures 3 and 4. **(A)** Genomic blot to detect telomere restriction fragments (TRF) in senescent WI38 and MRC5 cells grown at 20 and 3% oxygen. **(B)** USTELA assays on MRC5 cells grown to senescence at 20% or 3% oxygen as indicated. **(C)** Quantification of short USTELA products as detected in (B). The fraction of USTELA products running at or below 1.8 kb was determined. Each data point represents one PCR reaction. **(D)** Quantification of the number of γ-H2AX foci per nucleus in proliferating MRC5 cells treated for 2.5 h with zeocin (50 µg/mL) or etoposide (2.5 mM). n≥3 with 20-25 cells scored in each independent experiment. **(E)** Western blot for HIF-1α showing knockdown by an inducible shRNA to HIF-1α in proliferating MRC5 cells grown at 3% oxygen. **(F)** IF for HIF-1α in cells as in (E). **(G)** Quantification of the effect of HIF-1α depletion on zeocin-induced γ-H2AX foci in proliferating MRC5 cells grown at 20% and 3% oxygen. **(H)** Growth curves of MRC5 cells grown at 3% oxygen and treated with ATMi1, HIF-1αi, or both. ATMi1, KU-60019 at 670 nM; HIF-1αi, KC7F2 at 20 µM. The inhibitors were added on day 80. **(I)** Growth curves of MRC5 cells grown at 3% oxygen treated with ATMi1 (670 nM) and/or NAC (5 mM) as indicated. Compounds added at day 87. **(J)** Quantification of ROS and Superoxide in proliferating MRC5 cells grown at 3% and 20% oxygen. Assay as in Fig. 4A and B. **(K)** As in (J) but for WI38 cells. **(L)** Quantification of ROS species in proliferating MRC5 cells grown at 3% or 20% oxygen expressing Myc-TRF2 or the empty vector control. Assay as Fig. 4A and B. **(M)** Quantification of γ-H2AX foci in RPE1 cells treated with zeocin (50 µg/mL, 2.5 h), TCEP (0.5 mM, 2 days), or both. n = 2 with 20 cells per experiment. **(N)** Growth curve of MRC5 cells grown at 3% oxygen and treated with ATMi1, TCEP, or both. ATMi1, KU-60019 at 670 nM; TCEP at 0.5 mM. P values were determined as follows: (C), (J) and (K), unpaired t-test; (D) and (G) Ordinary One-Way ANOVA with Šídák’s multiple comparisons test; (L) Brown-Forsythe and Welch ANOVA tests with Dunnet’s multiple comparisons test; (M) Ordinary One-Way ANOVA with Tukey’s multiple comparison test.

## STAR Methods

### METHOD DETAILS

#### Cell Culture

MRC5 (male) and WI38 cells (female) were obtained from the ATCC and cultured in EMEM + L-Glutamine (ATCC, 30-2003), 15% fetal bovine serum (FBS, Gibco, 16000-044), 1% penicillin/streptomycin (P/S, Gibco 15140-122). RPE1 p53^-/-^/Rb^-/-^ ^62^ (female), T2p1^41^ (female), HEK293FT (female) (from Thermofisher), U2OS^48^ (female), and Phoenix A (female)(from the ATCC) cells were grown in DMEM/F12 (1:1) (Gibco, 11330-032), 15% FBS, and 1% P/S. U2OS cells were supplemented with doxycycline (1 µg/mL) to induce ATM expression. All cells were grown at 37°C, 5% CO2 in a humidified incubator (Thermo Heracell 240, Cat # 51019560). Cells were grown at normoxia or in an incubator maintained at 3% oxygen (Binder, Model CB170, Cat # 9140-0140). Media was replaced every 3-5 days, depending on the proliferation of cells. Time spent outside of 3% oxygen incubator was minimized. WI38 and MRC5 cells were trypsinized (Gibco, 25200-056) for ∼3-5 min at 37°C, in the oxygen condition-relevant cell incubator. Cells were counted at each passage (Beckman Coulter Z1 Particle Counter Dual, Cat # 6605699) to calculate population doublings using the standard equation: Log(Total/Seeded)*3.32.

#### Inhibitors

Inhibitors used were as follows: ATMi/ATMi1, KU-60019 (Selleckchem S1570, DMSO); ATMi2, AZD1390 (Selleckchem S8680, DMSO); ATMi3, KU-55933 (Selleckchem S1092, DMSO); ATRi/ATRi1, AZD6738 (Selleckchem S7693, DMSO); ATRi2, AZ20 (Selleckchem S7050, DMSO); ATRi3, M4344 (Selleckchem S9639, DMSO); Chk2i, BML-277 (Selleckchem S8632, DMSO); Chk1i, CHIR-124 (Selleckchem S2683, DMSO). Inhibitors were dissolved and diluted in 0.05% DMSO. Media with inhibitors or other compounds was changed every 3-4 days. For NAC treatment a 0.5 M solution (filter sterilized) of N-acetyl Cysteine (NAC) (Thermofisher, A15409.14) was used to add NAC at 5 mM to culture media buffered with HEPES-NaOH (5 mM, pH 7.5). NAC media was replaced every 3-4 days. For γ-H2AX IF, cells were incubated with 5 mM NAC for 2 days before induction of DNA damage.

#### BrdU incorporation assay

Approximately 40,000 and 15,500 cells were plated in 35 mm or 24-well microscope-compatible dishes, respectively (Ibidi, 81156 and 82426). After 3 days, media was replaced with media containing 10 µM BrdU (Sigma, B5002) together with relevant inhibitors in 0.05% DMSO. Cells were incubated for 4 days, washed with PBS (Corning, 21-040-CV), and fixed in 2% formaldehyde. After two more PBS washes, β-gal staining was performed using CellEvent Senescence Green Detection Kit (Invitrogen, C10851) according to manufacturer instructions. 4N HCl for 10 min was used to denature the DNA in preparation for BrdU staining. After three PBS washes, cells were blocked for 1 h using IF blocking buffer (1 mg/mL BSA, 3% Goat Serum, 0.1% Triton-X 100, 1 mM EDTA pH 8.0), followed by 1 h of anti-BrdU Alexa647-conjugated antibody (Abcam, ab220075) at 1:500 in IF blocking buffer combined with anti-Laminb Coralite594-conjugated antibody (Proteintech, CL594-66095) at 1:250-1:500. Samples were kept in the dark during and after BrdU/LaminB1 Ab incubation. After antibody incubation, samples were washed once with 1X PBS, then incubated in DAPI (0.17 µg/mL in 1X PBS) for five min, followed by another 1X PBS wash. Samples were then incubated in 2% formaldehyde for 10 min, followed by three 1X PBS washes. Fixed samples were imaged on the CellDiscoverer 7 (Zeiss) at 5X or 10X magnification, with the following LED settings: DAPI 385 nm 75% gain, 190 ms exposure at 30% intensity; AF647 at 625 nm 50% gain, 1400 ms exposure at 30% intensity; AF488 470 nm 100% gain, 150 ms exposure at 30% intensity; AF594 590 nm 75% gain, 200 ms exposure at 30% intensity. Images were then quantified based on co-localization of DAPI with BrdU/LaminB/β-gal IF signal using FIJI/ImageJ. Brightness/contrast settings were adjusted to minimize background fluorescence (as judged by fluorescent signal appearing outside the nucleus) using the same settings for all tested samples in each experiment.

#### Immunofluorescence

IF for γ-H2AX foci was performed on cells plated on PolyL-Lysine-coated coverslips (Corning, 354085) in 6 well dishes (Falcon, 353046) three days before drug treatment. NAC (5 mM final) was added at least two days before drug treatment and replenished during drug treatment. To induce DNA damage, zeocin, etoposide, or HU-containing media was equilibrated at 20 or 3% oxygen for ∼2.5 h before addition to the cells. Cells were washed with PBS equilibrated at 3% oxygen for >1.5 h and fixed in 2% formaldehyde in 3% oxygen-equilibrated PBS. Samples fixed for 5 min at 3% oxygen, followed by 5 min at 20% oxygen. All samples from cells grown at 20% oxygen followed the same procedure but with atmospheric oxygen conditions. Cells were subsequently treated as above (BrdU incorporation assay) except that there was no HCl denaturation. γ-H2AX antibody (Millipore, 05-636) was used at 1:2000 in IF blocking buffer. The secondary antibody was used at 1:500-1:1000 dilution (Alexa488, Invitrogen A11070). After the final fixation step, coverslips were mounted on microscope slides (Fisher, 22042944) with antifade gold (Invitrogen, P36934) and sealed with nail polish hardener (Sally Hansen). Slides were imaged at 60X on a DeltaVision microscope (Applied Precision microscope with Coolsnap QE Photometrics camera and SoftWoRx software). Foci were counted manually using ImageJ/FIJI. HIF-1α IF was performed as above with an anti-HIF-1α antibody (Abcam, ab179483) diluted at 1:500 – 1:1000. U2OS cells were also imaged at 100X on a Leica DMI-8 inverted microscope with a K8 cmos camera running THUNDER software.

#### IF-FISH for telomere damage

IF-FISH to measure the TIF response in T2p1 cells was performed as described^63^. TIFs were imaged at 60X or 100X on a Deltavision microscope and scored manually using ImageJ/FIJI.

#### Western blotting

Western blotting was performed as described^64^. Cell lysates were prepared with RIPA Lysis buffer with 1X Phos-Stop (Roche, 4906837001) and 1X complete protease inhibitor tablets (Roche, 11836170001). 100 mL RIPA (Thermofisher, 89900) was used for 500,000 cells, after which LDS was added to 1x (from a 4x stock, Thermofisher, NP0007) and βME was added to 350 mM (1:10 βME to LDS). Samples were boiled for 5 min and loaded into SDS-PAGE gels (Invitrogen, XP04122BOX or NP0322BOX). Western blot membranes were blocked in 5% BSA in TBST (TBS + 0.5% Tween20). Primary antibody incubations were done overnight at 4°C or 1 h at room temperature. After 3 washes (3 min each) in TBST, membranes were incubated for 1 h at room temperature, followed by 3 more washes in TBST (3 min each). Blots were developed with chemiluminescence reagent Supersignal West Pico PLUS (Thermofisher, 34580). Antibodies: TRF2, #647 ^65^; TRF1, #371 ^66^; Rap1, #765 ^64^ ; TIN2, #864 ^67^; TPP1, #1150^64^; POT1, #978 ^68^ or Proteintech 10581-1-AP; Vinculin, CST 13901; GAPDH, Invitrogen MA5-15738; Mre11, CST 4895; Chk2, CST 2662; HIF-1α, Abcam ab179483; ATM, CST 2873; ATM (For RPE1 and U2OS, Fig. 4), Sigma, A1106 1:7500. γ-tubilin GTU-88, Sigma T6557.

#### Analysis of ATM on non-denaturing gels

Protocol performed as described elsewhere ^57^. Briefly, cells were grown in relevant oxygen conditions for at least 72 hours. Cells were treated with NAC for 48 hours. Cells were washed with oxygen-acclimated PBS, then treated with 100 mM N-ethylmaleimide (NEM) (Thermofisher, 23030) in PBS for 5 minutes at 37°C in appropriate oxygen condition. Cells were then scraped and left to sit on ice for 15 minutes. After spinning cells down (5 mins, 4000xg, 4°C), pellet was resuspended in 250 uL NP-40 non-denaturing lysis buffer (100 mM NaCl, 5 mM CaCl_2_, 20 mM Tris-Cl pH 7.5, 0.5 % (v/v) Nonidet P-40 (NP-40),) and incubated for 15 mins on ice, infrequent vortexing. 1 uL Benzonase Nuclease (Millipore, E1014) was added to each sample and put at 37°C for 10 minutes. 0.5M EGTA was added at 1 uL/100 uL lysis buffer to stop the reaction. SDS was then added to a final concentration of 1%, and pellets were then incubated on ice for another 15 minutes. Tibes were then spun down at 14000xg for 10 minutes at 4°C, supernatants were transferred. Protein was measured, and 20 µg were loaded per lane on a 3 – 8% poly-acrylamide Tris-acetate native gel (NuPage,TA03810BOX); run at 65 volts until 250 kd protein marker was halfway down the gel, followed by overnight transfer at 30 volts, 4°C. Antibody for ATM was Sigma, A1106 1:7500. Lifting cells by trypsin instead of scraping did not seem to affect outcome of the assay.

#### Live-cell imaging

Cells were plated in 35 mm or 24-well Live-Cell Microscopy plates (Ibidi 81156 and 82426, respectively) and cultured for 2-3 days. Media was replaced with fresh MEM without phenol-red (Corning, 17-305-CV; supplemented with 1% sodium pyruvate (Sigma, S8636), 1% HEPES, 1% L-glutamine, and 15% FBS) containing drugs (or DMSO) on the day of imaging. Media for cells grown at 3% oxygen was pre-equilibrated at 3% oxygen for 1.5 h. Cells were imaged at 30, 45, or 60 min intervals depending on the experiment on a CellDiscover 7 microscope using ZEN3.6 software (Zeiss). Magnification was 5X or 10X, depending on the experiment. Channel intensities are as follows: PGC at 5% intensity, 6 ms exposure time with 70% intensity; mCherry LED 590nm at 75.2% intensity, 500 ms exposure at 30% intensity; TaYFP 5470 nm at 70.3% intensity, 190 ms exposure at 30% intensity. Imaging conditions: 5% CO2, 37°C, and atmospheric oxygen. When brightness/contrast changes during or after imaging were required, changes were made consistently for different conditions compared within each experiment. The first 7 h of each movie was deleted because of high background at the beginning of each imaging session.

#### Lentiviral and retroviral infections

Lentiviral infections were performed as described ^69^. One day before transfection, HEK293FT cells were plated at ∼4.5 million cells per 10 cm dish. Calcium phosphate transfection was used to introduce 15.4 µg of lentiviral construct DNA, 15.4 µg pPAX2, and 10 µg pMD2. 12-18 h after transfection, HEK293FT cells were switched to the media appropriate for the cells to be infected, incubated for 24 h, and the media was filtered through a 0.4 µm filter. After addition of polybrene (4 µg/mL) the filtered media was added to target cells. Media on HEK293FT cells was replaced, allowing for 2-3 more applications of filtered media as required. Lentiviral plasmids: tFucci(CA)2, RIKEN BioResource Center, RDB15446; shHIF-1α, plKO.1-TetON-Puro-Hif1a 0819 (Addgene 118704).

Retroviral infections were performed as described ^64^. Briefly, Phoenix A cells (∼4.5-5 million) were plated in 10 cm dishes one day before transfection. Calcium phosphate transfection was used to introduce 20 µg retroviral construct DNA and 0.5 µg of a GFP expression plasmid as a control for transfection efficiency. At 12-18 h after transfection, media was changed to that of target cells and infection was performed as above for lentiviral infections. Expression constructs: pLPC-Myc-TRF2 ^65^; pLPC-FLAG-Rap1 ^71^; pLPC-TPP1-IRES-FLAG-POT1 was generated by Gibson cloning using pLPC-FLAG-POT1 ^63^ and geneblocks (IDT).

#### USTELA

USTELA was carried out as previously described^40^. Briefly, cells were trypsinized, collected (5,000 <u>g</u>, 5 min, 4°C) and flash frozen as pellets or processed immediately. Genomic DNA (gDNA) was isolated using the MagAttract HMW Kit (Qiagen, 67563). In our hands, phenol-chloroform extraction of gDNA was not compatible with USTELA. DNA was digested overnight with *Mse*I (60 U, NEB, R0525L) and *Nde*I (60U, NEB, R011S). Digested DNA (20 ng) was ligated to the 42-mer and 11-mer as described previously^40^. After ligation, 1 nM telorette mix and 1 mL 1:10 diluted T4 DNA Ligase was added, alongside T4 Ligase Buffer and ddH20 and incubated overnight at 35°C. The next day, samples were heat-inactivated for 20 min at 65°C and diluted to ∼5 pg/mL. PCR was performed with Failsafe PCR 2X Mastermix (Lucigen, FSP995H), Failsafe Enzyme (Lucigen, FS99060), and Tel-tail and Adapter primers in 24 µL final volume. PCR conditions: 68°C 5 mins (1X), 95°C 2 mins(1X), 95°C 15s, 58°C 30s, 72°C 12 mins (26X), 72°C 15 mins (1X). Samples were run on a 1% agarose gel in 1XTAE. Gels were blotted and hybridized with a TTAGGG repeat probe as described^69^.

#### Genomic blots to determine TRF length

Southern blotting to measure the length of telomeric *Mbo*I/*Alu*I fragments was performed as previously described ^72^.

#### Comet assay

Comet assays were performed using the CometAssay Kit (Biotechne R&D Systems, 4250-050-K) with modifications to avoid changes in zeocin-induced DSBs during the procedure. Cells were treated with 50 µg/mL zeocin for 2 h. Zeocin containing media used for low oxygen was equilibrated to 3% oxygen. After treatment, cells were harvested with trypsin, spun down, and resuspended to a density of ∼200,000 cells/mL in PBS equilibrated at 3% oxygen. 5 µL of diluted cells was added to 45 µL melted CometAssay LMAgarose (R&D systems, 4250-050-02), mixed, and spread on comet assay slides. Slides were left at 3% or 20% for 30 min and then for 40 min at 4°C in 20% oxygen. Slides were covered with lysis solution for 25 min at 37°C in 3% or 20% oxygen, followed by 35 min at room temperature at 20% oxygen. Slides were processed according to the manufacturer’s instructions and analyzed using the publicly available OpenComet ImageJ plug-in^73^ (https://cometbio.org).

#### Tracking of cell cycle stages in live-cell imaging

Raw data generated from TrackMate was visualized on individual-cell and full-experiment scales using a custom Rscript. Because TrackMate requires a designated channel in which to look for trackable units, we created a new channel by combining the RFP and YFP images. Frames with errors in auto-tracking were manually added or removed within TrackMate. A “background track” was manually generated to control for noise and background in the written code by tracking a space in the imaging field with no cells throughout the video. This track was always the last track made (with the highest TRACK ID in Trackmate) for the Rscript analysis.

The major steps the Rscript takes are as follows (see also Figure S2B). All conditions are first combined into a single data table, on which the 5^th^ percentile value (maximum value of the bottom 5%) for red (mCherry) and green (mVenus) fluorescence are determined. The 5^th^ percentile serves as a baseline that defines “background,” and for both channels, each value above background is converted to a “fluorscore,” or its number of standard deviations above the baseline. The fluorscores are then passed through a “smoothing” (local regression analysis) “loess” function to neutralize spurious fluorescence spikes or drops caused by cells entering and exiting focus (Figure S2B). Three different intensities of smoothing are applied and manually checked, ensuring there is no over-correction leading to flattened fluorescence curves. Two other modifiable parameters in the form of “general background correction” are applied for mCherry and mVenus (HBRc and HBRv, respectively). HBRc and HBRv are numerical constants (0.5-3) that are applied to the background fluorescence, fluorscore, and z-score values defined by the “background track”. JPEG image outputs for all individual cell tracks are generated with each output track containing six traces: raw fluorescence, z score, fluorscore, small smoothed fluorscore, medium smoothed fluorscore, and large smoothed fluorscore. The x-axis represents the movie frame from which the graphed value came. Values of mCherry and mVenus are shown as separate red and green lines. HBRc/v altered background values are denoted via horizonal dashed lines (see Figure S2B).

For global analysis, we used the fluorscore values to predict the cell cycle phase of a cell in any given frame by passing the data through two functions. The first defined the cell cycle phase (G1, S, or G2) by comparing mCherry and mVenus fluorscore values to their baseline and to each other. The full list of parameters is available at https://github.com/astu17/Trackmate-FUCCI-Code.git. The second function corrects anomalous, aberrant calls that the first function may have made. Each cell track is mapped on a graph where x is the tracked frame, and y is a different cell track. Each block in a track is colored according to its associated cell cycle phase. Tracks are sorted on the y-axis based on the amount of cell cycle-active frames. The x-axis origin represents the beginning of a given cell track. In all cases, each track was manually checked to confirm the given output. Erroneous outputs were manually corrected. Four visualizations of the data are generated: one based on the raw fluorscore, and one for each level of fluorscore smoothing.

In the final tracks presented, G1 cells are gray and S and G2 cells are presented into a single green color indicating cell cycle activity. Divisions were manually labelled with a yellow bar at the frame of division. High initial background necessitated beginning our tracking ∼7 h after imaging started. Some videos had more cells than were quantified, and cells tracked were picked at random. In some cases, cells that remained in G1 were incorporated into total quantification while not being included in the tracking.

#### Detection of oxidative stress

ROS and Superoxide were detected using a ROS/Superoxide Detection Kit (abcam ab139476) according to the manufacturer’s instructions with slight modifications. Briefly, cells were plated in 35 mm microscope-compatible plates (Ibidi, 81156) and allowed to settle in 20% or 3% oxygen for 3 days. On the third day, reagents to detect ROS and superoxide were added to fresh media, which was then acclimated to the appropriate oxygen condition (20% or 3%, 1 hour in the respective incubator). Cells were then incubated for one hour with matched media containing detector reagents. After 30 minutes, 50 µM pyocyanin was added to a 20% plate as a positive control. After washing as per the manufacturer’s instructions, cells were quickly imaged on a CellDiscover 7 microscope using ZEN3.6 software (Zeiss). Magnification was 20X. Nuclear intensities quantified using channel intensities determined via TrackMate.

#### Cysteine Reactivity Mass Spectrometry

Cysteine reactivity profiling was performed following a modified protocol from Vinogradova et al. 2020^56^.Cells were washed with PBS that had been acclimated to the appropriate oxygen condition, lifted either by scraping in cold PBS or by trypsinization, spun down at 4,000 *g* and flash frozen as pellets and kept at -80°C until processing. Pellets were then resuspended in 560 µL PBS and lysed by sonication (3 x 8 pulses, on ice, 60%). Protein concentration was then measured via BCA assay (Pierce, 23225), and samples were diluted in PBS to a protein concentration of 1 mg/mL. All tubes used in this workflow were low-binding tubes (Eppendorf, 022431081). 5 µL of 10 mM iodoacetamide polyethyleneoxide desthiobiotin (IA-DTB, Santa Cruz) was added to each sample and mixed gently (no vortexing). After 1h incubation at room temperature with rotation, samples were transferred on ice and proteins were precipitated using MeOH/CHCl_3_ precipitation. Cold MeOH (500 µL) followed by CHCl₃ (100 µL) were added to each tube, and proteins were pelleted by centrifugation at 10,000 × g for 10 min at 4°C. Tubes were then vortexed and spun down at 10,000 <u>g</u>, 10 min, 4°C. Supernatant was aspirated to isolate the resulting protein disk. An additional 400 µL MeOH was added, samples were sonicated to break up the pellet, and proteins were pelleted by spinning at 10000 <u>g</u>, 10 min, 4°C. The pellets were resuspended in 90 µL of 9M urea, 10 mM DTT, and 50 mM triethylammonium bicarbonate buffer (pH 8.5) mixture. Samples were heated at 65°C for 20 min, followed by cooling to room temperature and addition of 10 µL of 500 mM iodoacetamide in H_2_O. Samples were then agitated at 37°C for 30 min in the dark. 300 µL of 50 mM TEAB (pH 8.5) was added to reduce the UREA concentration, and samples were sonicated to solubilize proteins. 4 µL of 100 mM CaCl_2_ was added, followed by addition of 4 µl of 0.5 µg/mL trypsin/LysC (Promega). The digestion took place overnight at 37°C with shaking. The next day, samples were centrifuged to sediment undigested material and the supernatant was transferred to a new tube, followed by addition of pre-washed streptavidin-agarose beads (Fisher, 20349) to each sample for enrichment (50 μL beads, resuspended in 50 mM TEAB buffer, pH 8.5, containing 150 mM NaCl and 0.2% NP40; 300 μL/sample) and the mixture was rotated for 3 h at room temperature. Following the enrichment step, the beads were pelleted, transferred to BioSpin columns and washed (3 x 1 mL of wash buffer (50 mM TEAB buffer, pH 8.5, containing 150 mM NaCl and 0.1% NP40), 3 x 1 mL PBS, 3 x 1 mL H_2_O). The peptides were eluted into a new Eppendorf tube by adding 300 μL of 50% HPLC-grade acetonitrile, containing 0.1% FA. The eluate was removed by SpeedVac vacuum concentrator. TMT labeling, HPLC fractionation and Mass spectrometry analysis were then performed as previously described^56^.

#### Data processing and analysis for cysteine reactivity profiling

MS1, MS2, and MS3 files were extracted from RAW files using RawConverter (available at https://github.com/proteomicsyates/RawConverter) and searched using the ProLuCID algorithm with a reverse concatenated version of the Human UniProt database (release 2024-03), using the Integrated Proteomics Pipeline (IP2). Peptides were searched with up to one differential modification for the IA-DTB probe (+398.25292 Da) and a static modification for carboxyamidomethylation (+57.02146 Da). An additional static modification for the TMT-tag (+304.2071 Da for the 16-plex experiments or +229.162932 Da for the 6-plex experiment) was set for lysine residues and N-termini. ProLuCID data was filtered through DTASelect to achieve a peptide-false positive rate below 1%. The reporter ion mass tolerance was set to 20 for MS3-based peptide quantification in IP2.

Each experiment was processed individually with the requirements: removal of half-tryptic peptides, removal of peptides with more than two internal missed cleavage sites, removal of peptides with <5000 average reporter ion intensity across all technical replicate groups and removal of peptides with high variation between all technical replicate groups (coefficient of variation > 0.5). Peptide ratios (signal intensity/sum of signal intensities per peptide) were calculated and multiplied by the number of channels to account for differences in multiplexing between experiments. Peptide ratios were averaged (median) for multiple cysteines on the same peptide and multiple peptides containing the same cysteine. Peptides were required to be quantified in at least 2 experiments to be included in the final table.

## QUANTIFICATION AND STATISTICAL ANALYSIS

Error bars represent mean ± standard deviation unless otherwise stated. The number of replicates and statistical tests used are indicated in the figures and figure legends. Statistics calculated using Graphpad Prism 10.2.

## ADDITIONAL RESOURCES

Rscript code is available online at https://github.com/astu17/Trackmate-FUCCI-Code.git.

## Key resources table

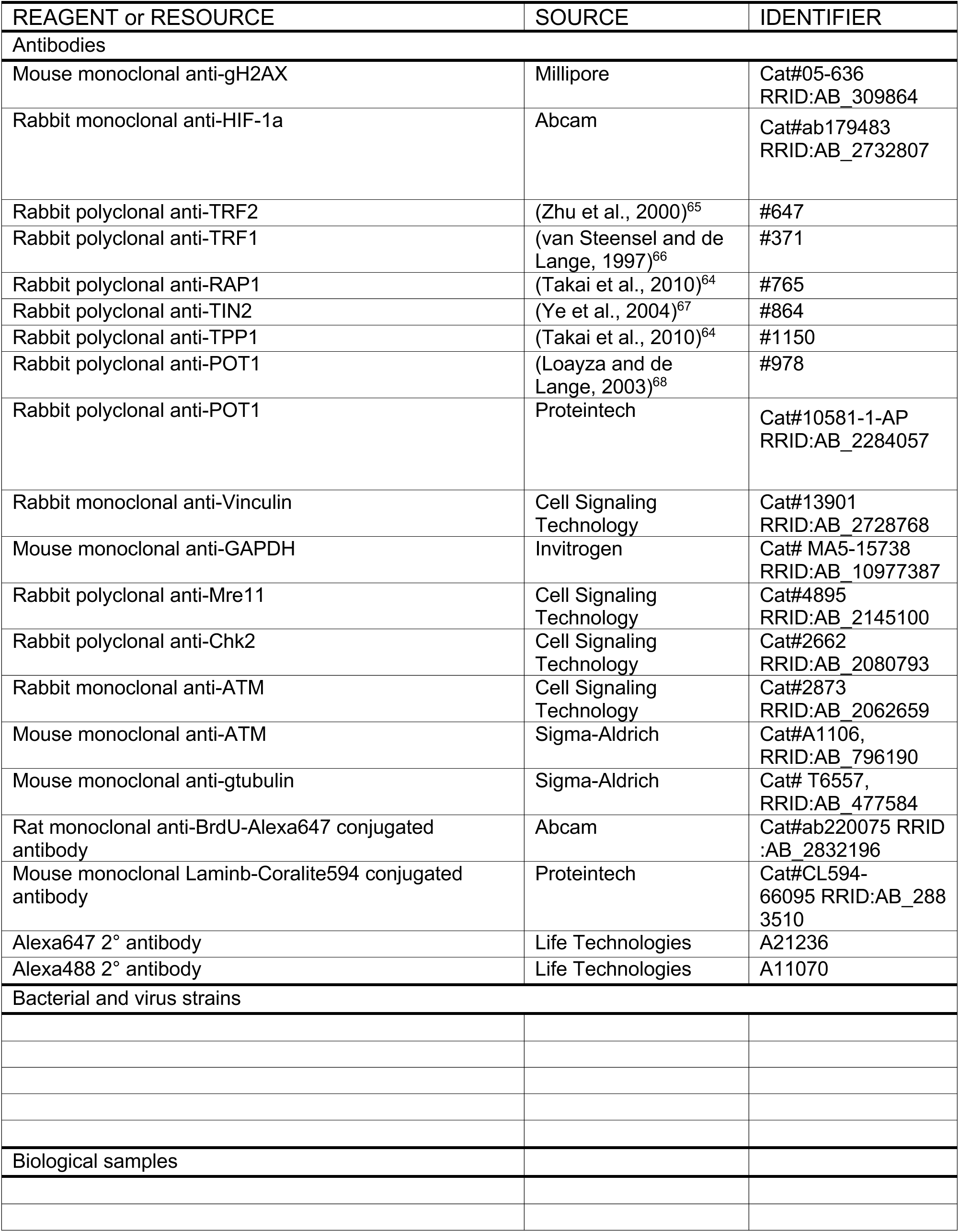
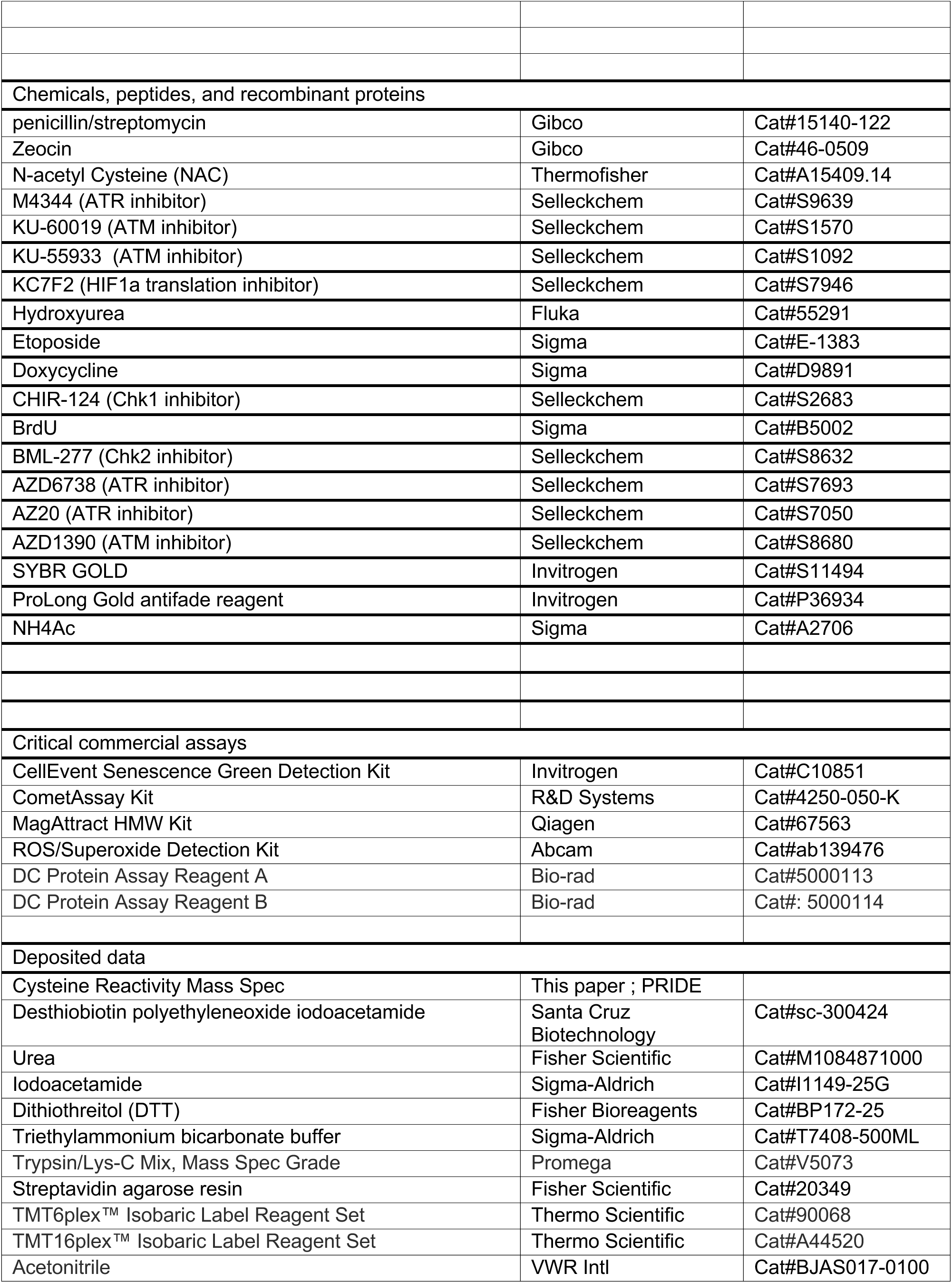
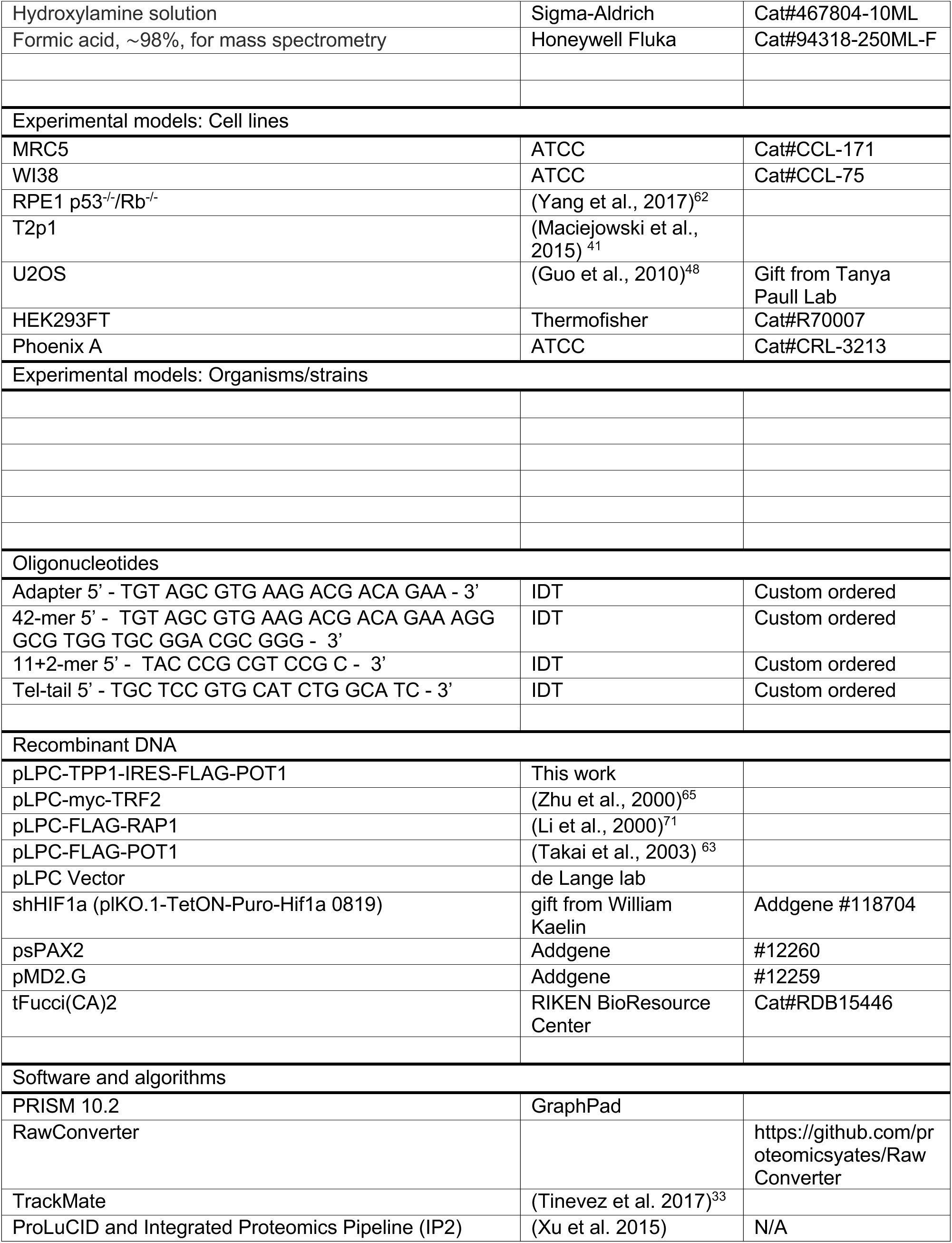
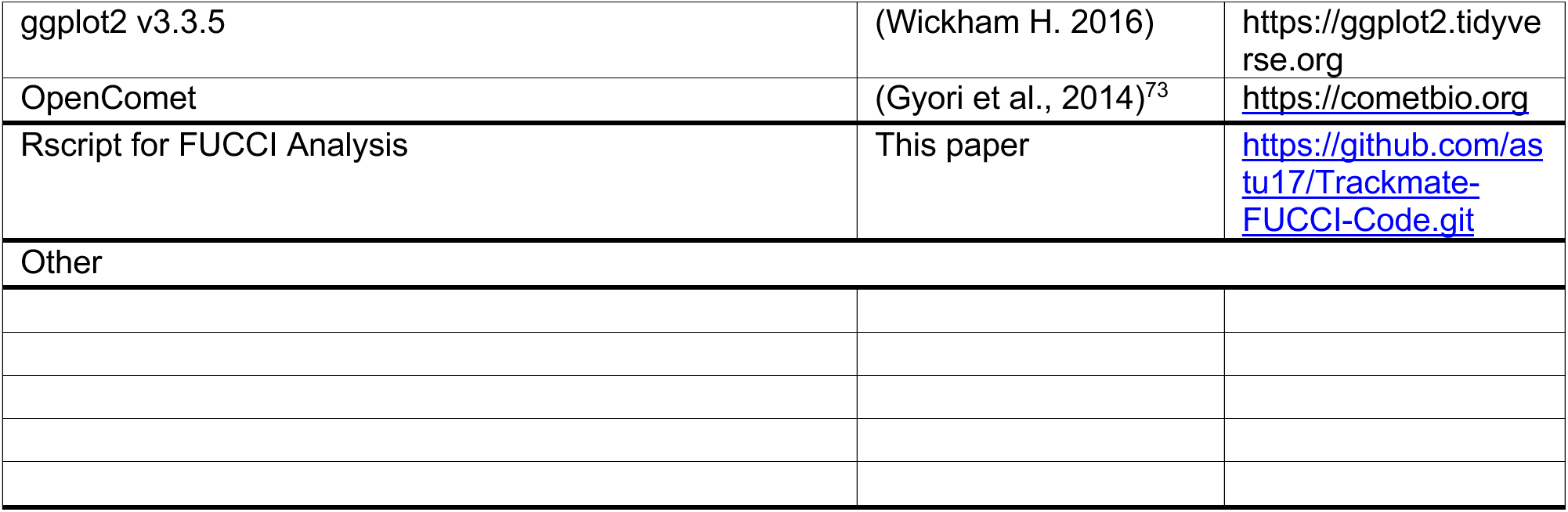

## Supplemental Item Titles

**Supplementary Video S1** Example of pre-senescent proliferating MRC5 cell.

**Supplementary Video S2** Example of MRC5 cell undergoing aberrant cell cycle event.

**Supplementary Video S3** MRC5 cell in a senescent population undergoing 3 divisions after ATMi treatment.

**Supplementary Video S4** MRC5 cell in a senescent population undergoing 2 divisions and 1 aberrant event after ATMi treatment.

**Supplementary Table S1.** Proteomics data of relevant cysteines from mass spectrometry / cysteine reactivity analysis.

